# Context-dependent sensory modulation underlies Bayesian vocal sequence perception

**DOI:** 10.1101/2022.04.14.488412

**Authors:** Tim Sainburg, Trevor S McPherson, Ezequiel M. Arneodo, Srihita Rudraraju, Michael Turvey, Brad Thielman, Pablo Tostado Marcos, Marvin Thielk, Timothy Q Gentner

## Abstract

Vocal communication in both songbirds and humans relies on categorical perception of smoothly varying acoustic spaces. Vocal perception can be biased by expectation and context, but the mechanisms of this bias are not well understood. We developed a behavioral task in which songbirds, European starlings, are trained to to classify smoothly varying song syllables in the context of predictive syllable sequences. We find that syllable-sequence predictability biases perceptual categorization following a Bayesian model of probabilistic information integration. We then recorded from populations of neurons in the auditory forebrain while birds actively categorized song syllables, observing large proportions of neurons that track the smoothly varying natural feature space of syllable categories. We observe that predictive information in the syllable sequences dynamically modulates sensory neural representations. These results support a Bayesian model of perception where predictive information acts to dynamically reallocate sensory neural resources, sharpening acuity (i.e. the likelihood) in high-probability regions of stimulus space.

**One-Sentence Summary:** Predictive information in vocal sequences biases Bayesian categorical perception through rapid sensory reorganization.

**Graphical Abstract:** 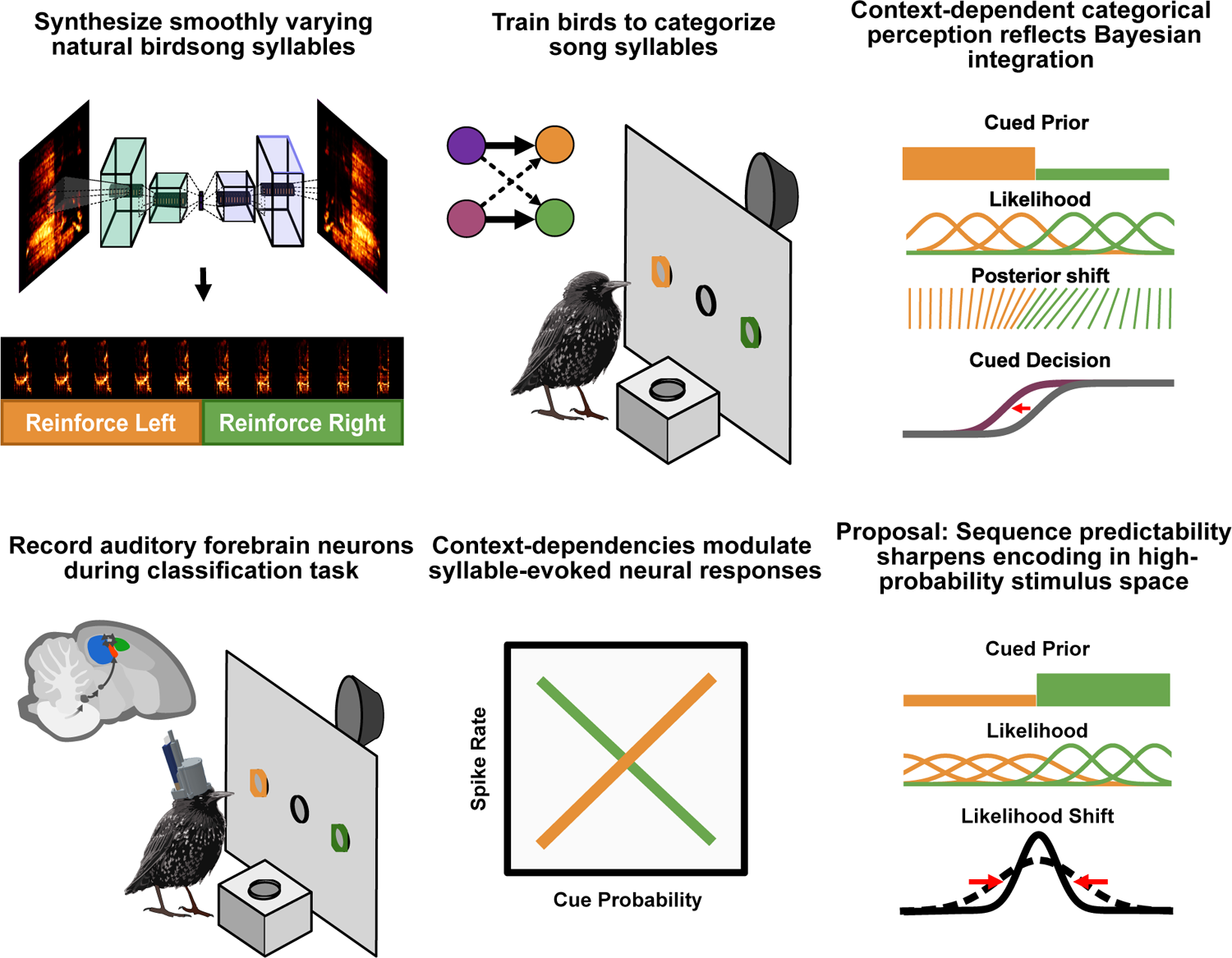

## 1 Introduction

Categorical perception, the grouping of smoothly varying signals into discrete classes, plays an important role in organizing complex experiences into shared representations by enabling abstraction and generalization across instances of explicit signals. Auditory categorical perception is fundamental to human vocal communication by speech. Categorical perceptual boundaries in speech are learned and differ across languages. Infants, for example, discriminate phonetic boundaries in speech that adults cannot (*1*). Phoneme perception is also influenced by the context of words, utterances, discourse, and environmental signals that bias perception towards more likely scenarios (*2–4*). For example, in the Ganong effect (*2*) categorical perception of ambiguous phonemes shifts based upon word expectations. The same ambiguous phoneme between ‘/b/’ and ‘/p/’ is more likely to be perceived as ‘peace’ than ‘beace’ but less likely to be perceived as ‘peef’ than ‘beef’, because ‘peace’ and ‘beef’ are common words. Context-dependent categorical perception in speech is also driven by the sequential organization of speech elements, including the relative positions of phonemes within words (*3*). The processes underlying categorical perception of phonemes in speech can be expressed as Bayesian inference over acoustic distributions (*4–6*). Under this framework, prediction bias, top-down influences, and prior expectations for speech perception are modeled as probabilistic integration. The specific cognitive and neural mechanisms underlying how predictive information and context-dependency modulate categorical perception of speech signals are not well-understood (*4*).

Categorical perception is not unique to human speech and has been observed in a number of sensory modalities and species (*1, 3, 7*). Songbirds perceive some elements of song categorically (*3, 8*), providing an opportunity to study mechanisms of categorical vocal perception in neurobiological detail. In swamp sparrows, somewhat similar to speech, categorical perceptual boundaries of notes are modulated by their position within the song (*9*). Birdsongs are learned and song repertoires can be maintained across populations for generations (*10*), suggesting that perceptual categories and the modulation of categorical perception may also be learned. A neural correlate for categorical perception has previously been described in the sensorimotor system of swamp sparrows (*8*), where spike rate differences in neurons in the auditory-motor nuclei HVC reflect natural vocal boundaries (*8*). It remains unknown for songbirds, how predictive information biases categorical perception and whether, as in speech, perceptual biases can be explained through Bayesian integration.

We developed methods to explicitly impose probabilistic predictive information in a sequence of birdsong syllables and trained European starlings, a songbird with complex vocal repertoires, to classify smoothly varying syllables while controlling sequential predictive song structure. We found that songbirds integrate learned sequential information in the categorical perception of vocal signals and that resulting perceptual biases are modulated by the strength of the sequential context and the uncertainty of the signal. We then explain how perceptual boundaries shift as a function of predictive information using a Bayesian account, successfully capturing all aspects of information integration, over the prior, likelihoods, and posterior probabilities.

We then explored the neurophysiological basis of context-dependent categorical perception using the same behavioral paradigm. We recorded spiking activity in behaving birds from populations of neurons in the primary auditory forebrain region Field L, two secondary auditory forebrain regions (the caudal mesopallium [CM] and caudal medial nidopallium [NCM]), and the caudal lateral nidopallium (NCL), a higher-order forebrain region implicated in visual and multi-modal working memory (*11–13*). We find that the sequential context of song syllables biases neural representations throughout these regions, leading to dynamic changes in sensory acuity (i.e. the likelihood of the Bayesian model) in predicted regions of acoustic space.

## 2 Results

### 2.1 Behavior

#### Paradigm

Sensory neuroscience and psychophysics have long, productive histories founded on the idea of relating parametric change in a stimulus to quantifiable changes in both neural activity (*14*) and behavior (*15*). Implicit in this approach is the strong assumption that sensory inputs can be discretized into stimulus events parametrically varying along one or two continuous dimensions. This approach is ideally suited to investigate how simple, non-natural, easily controllable, signals are perceived behaviorally or encoded neurally, but neglects the natural history of sensory systems adapted to ethologically relevant signals like birdsong (*16, 17*). Attempts to apply the same kind of parametric stimulus control to natural stimuli are rare, however, because natural signals tend to vary along multiple dimensions simultaneously (*18, 19*). We developed a behavioral paradigm to control context-dependent categorical perception in a natural stimulus space and capitalize on psychophysical methods by synthesizing smoothly varying starling song syllables using a neural network.

We captured the complex spectro-temporal statistics of song acoustics using a deep convolutional variational autoencoder (Fig 1A) (*20*) trained on a large library of conspecific song. From the latent space of this network, we synthesized acoustic continua (N=9), each comprising 128 synthetic syllables (morphs) that smoothly vary between two acoustically distinct syllable endpoints. We trained starlings using a two-alternative choice (2AC) category learning task to classify the naturalistic syllable morphs that lie along these continua (Fig 1B). We divided each continuum at the interpolation midpoint and reinforced one half with food for pecks into the left response port and other half for pecks into the right response port (Figure 1C bottom). We trained birds (n=20) on the syllable classification task to obtain psychometric functions for each syllable continuum (Fig 1D), then introduced cue syllables preceding the target (to-be-classified) syllable. Each cue syllable provided predictive information about the likely category of the target syllable (Fig 1E)). All subjects learned the task to at least 75% accuracy (Table 1) performing a total of 4.8 million behavioral trials.

**Figure 1:**
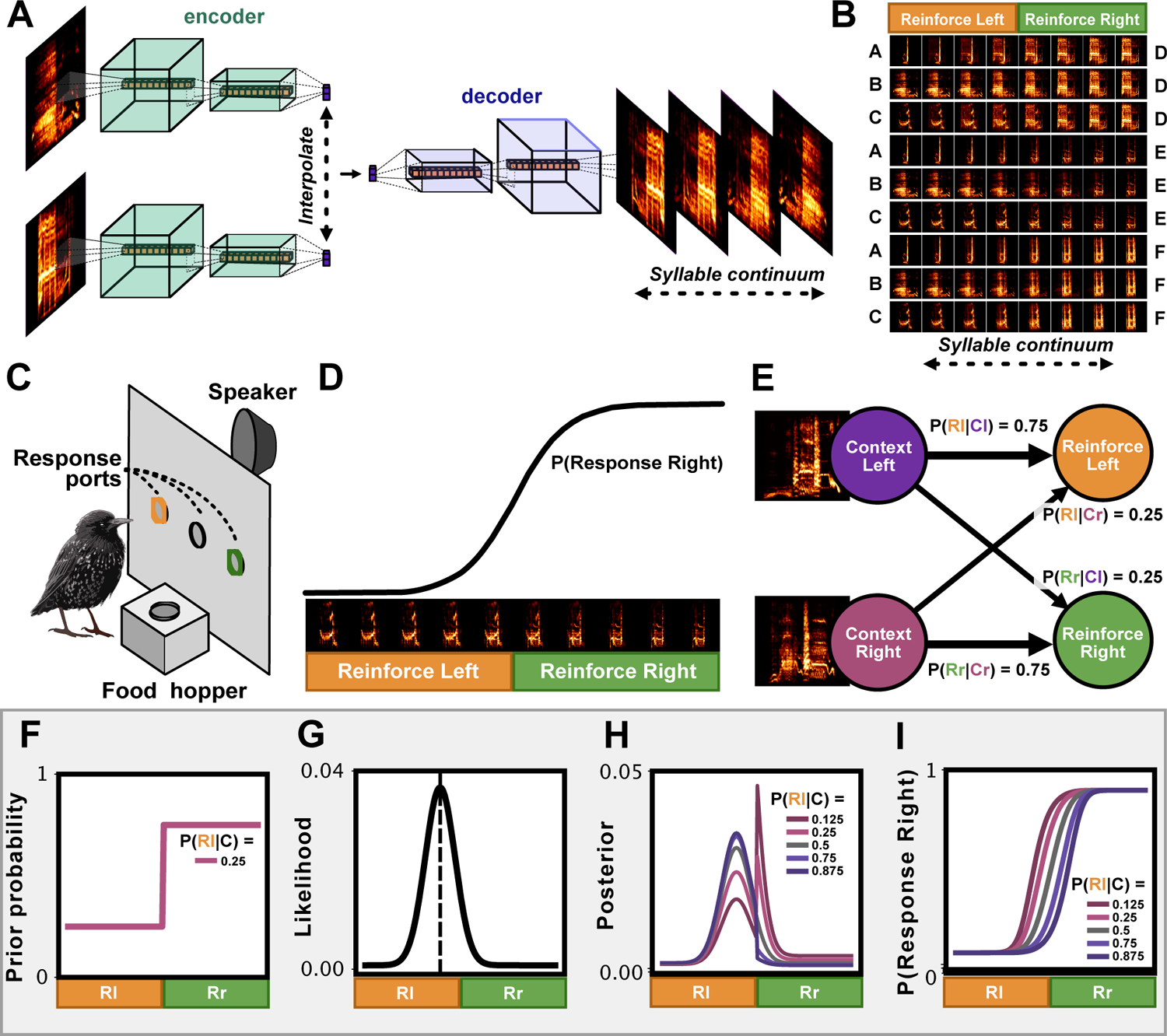
Behavioral assessment and model. (A) Syllable morphs are generated as interpolated projections between two song syllables in the latent space of a neural network. (B) Example syllables from the 9 morph continua (rows) used for behavioral training. The reinforced category is shown on the top, and the endpoint syllables are labeled on the left and right sides. (C) Operant apparatus used for this experiment. Green and orange response ports correspond to the syllable classes in (B). (D) A psychometric curve depicting syllable classification over one continuum. (E) Two example context cue syllables precede the target syllables, holding predictive information about the class to which each belongs. (F-I) A Bayesian model depicting our hypothesis. (F) An example prior probability represents the probability of a target syllable class given by the cue syllable (here the cue predicts a syllable associated with a right response). (G) Likelihood, given by a Gaussian distribution centered on the true target syllable (dotted line) on a given trial. (H) Posterior probability under the five cue (prior) probabilities used in this study. (I) The predicted behavior response under the Bayesian model, depicting a shift in categorical perceptual decision making as a function of the cue probability.

**Table 1:**
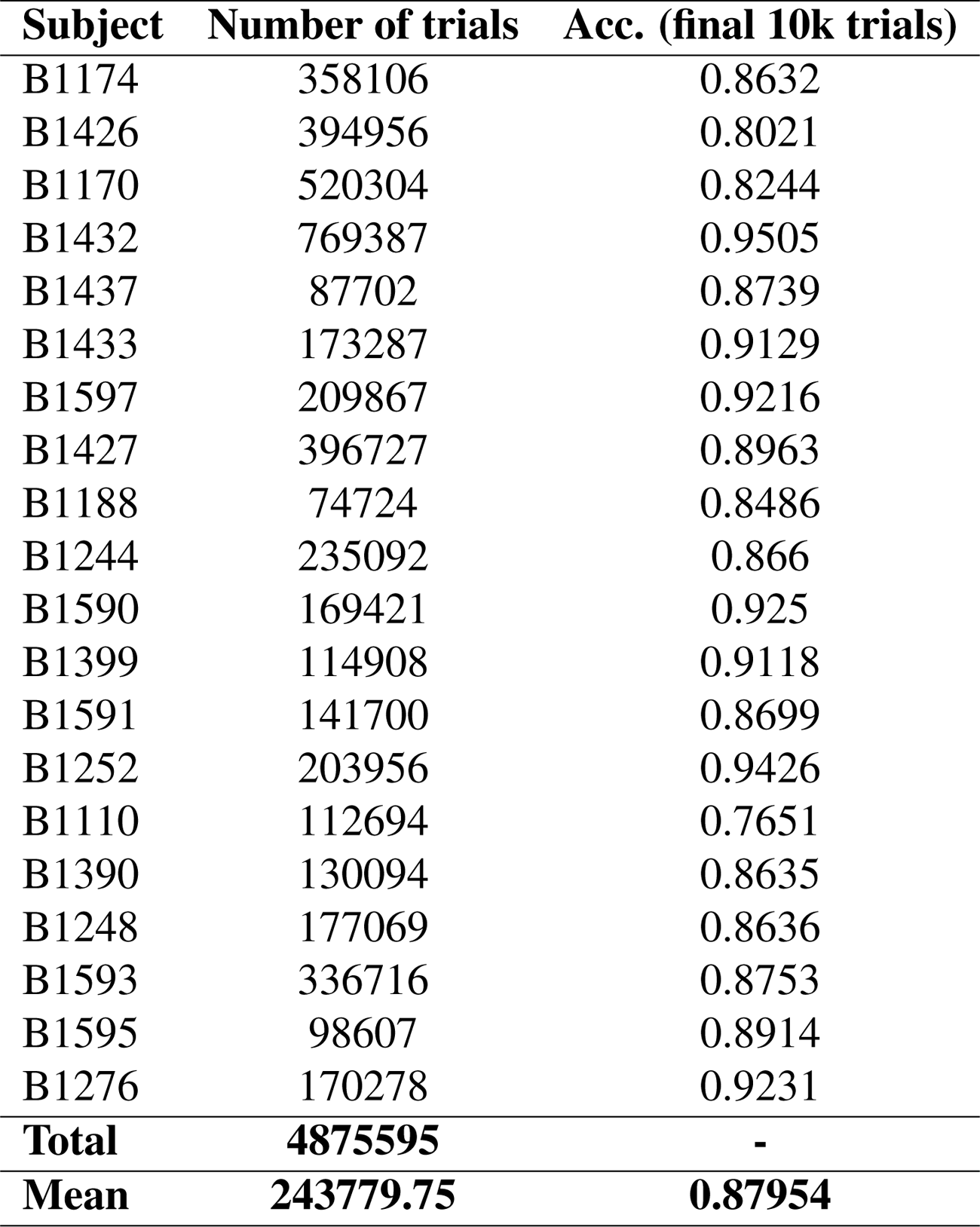
Behavioral datasets

| **Subject** | **Number of trials** | **Acc. (final 10k trials)** |
| --- | --- | --- |
| B1174 | 358106 | 0.8632 |
| B1426 | 394956 | 0.8021 |
| B1170 | 520304 | 0.8244 |
| B1432 | 769387 | 0.9505 |
| B1437 | 87702 | 0.8739 |
| B1433 | 173287 | 0.9129 |
| B1597 | 209867 | 0.9216 |
| B1427 | 396727 | 0.8963 |
| B1188 | 74724 | 0.8486 |
| B1244 | 235092 | 0.866 |
| B1590 | 169421 | 0.925 |
| B1399 | 114908 | 0.9118 |
| B1591 | 141700 | 0.8699 |
| B1252 | 203956 | 0.9426 |
| B1110 | 112694 | 0.7651 |
| B1390 | 130094 | 0.8635 |
| B1248 | 177069 | 0.8636 |
| B1593 | 336716 | 0.8753 |
| B1595 | 98607 | 0.8914 |
| B1276 | 170278 | 0.9231 |
| **Total** | **4875595** | **-** |
| **Mean** | **243779.75** | **0.87954** |

**Table 2:**
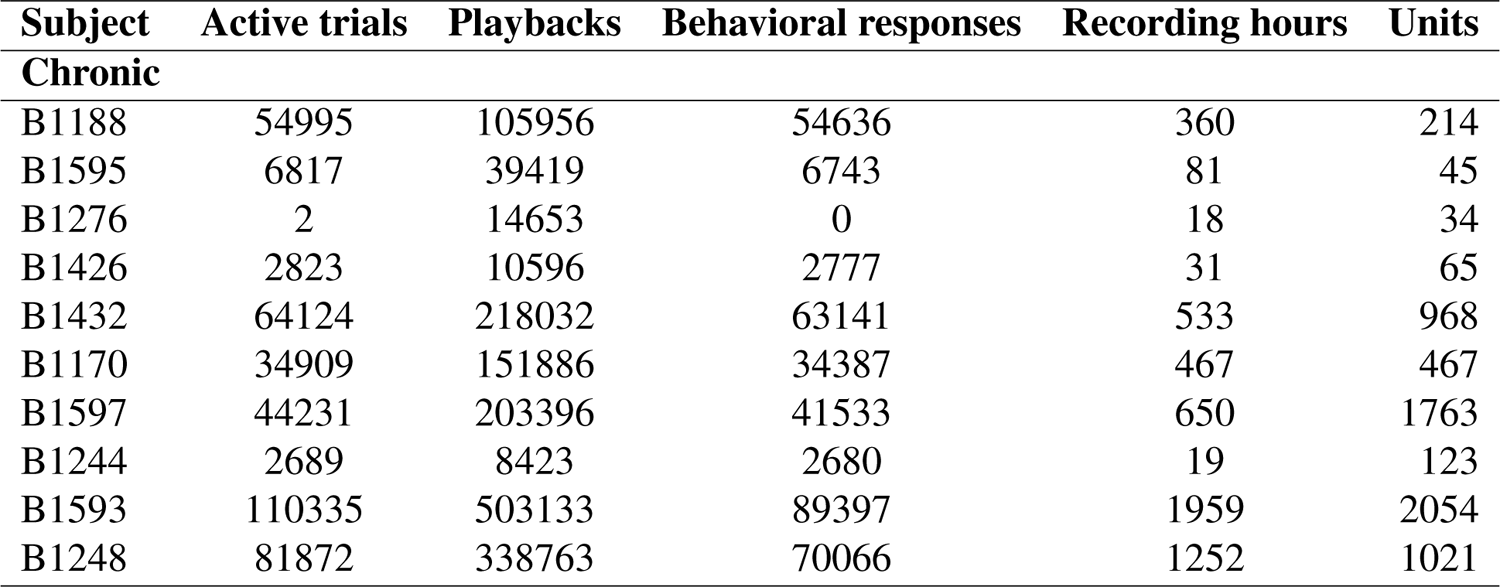
Neural datasets

| **Subject** | **Active trials** | **Playbacks** | **Behavioral responses** | **Recording hours** | **Units** |
| --- | --- | --- | --- | --- | --- |
| **Chronic** |  |  |  |  |  |
| B1188 | 54995 | 105956 | 54636 | 360 | 214 |
| B1595 | 6817 | 39419 | 6743 | 81 | 45 |
| B1276 | 2 | 14653 | 0 | 18 | 34 |
| B1426 | 2823 | 10596 | 2777 | 31 | 65 |
| B1432 | 64124 | 218032 | 63141 | 533 | 968 |
| B1170 | 34909 | 151886 | 34387 | 467 | 467 |
| B1597 | 44231 | 203396 | 41533 | 650 | 1763 |
| B1244 | 2689 | 8423 | 2680 | 19 | 123 |
| B1593 | 110335 | 503133 | 89397 | 1959 | 2054 |
| B1248 | 81872 | 338763 | 70066 | 1252 | 1021 |

**Table 3:**
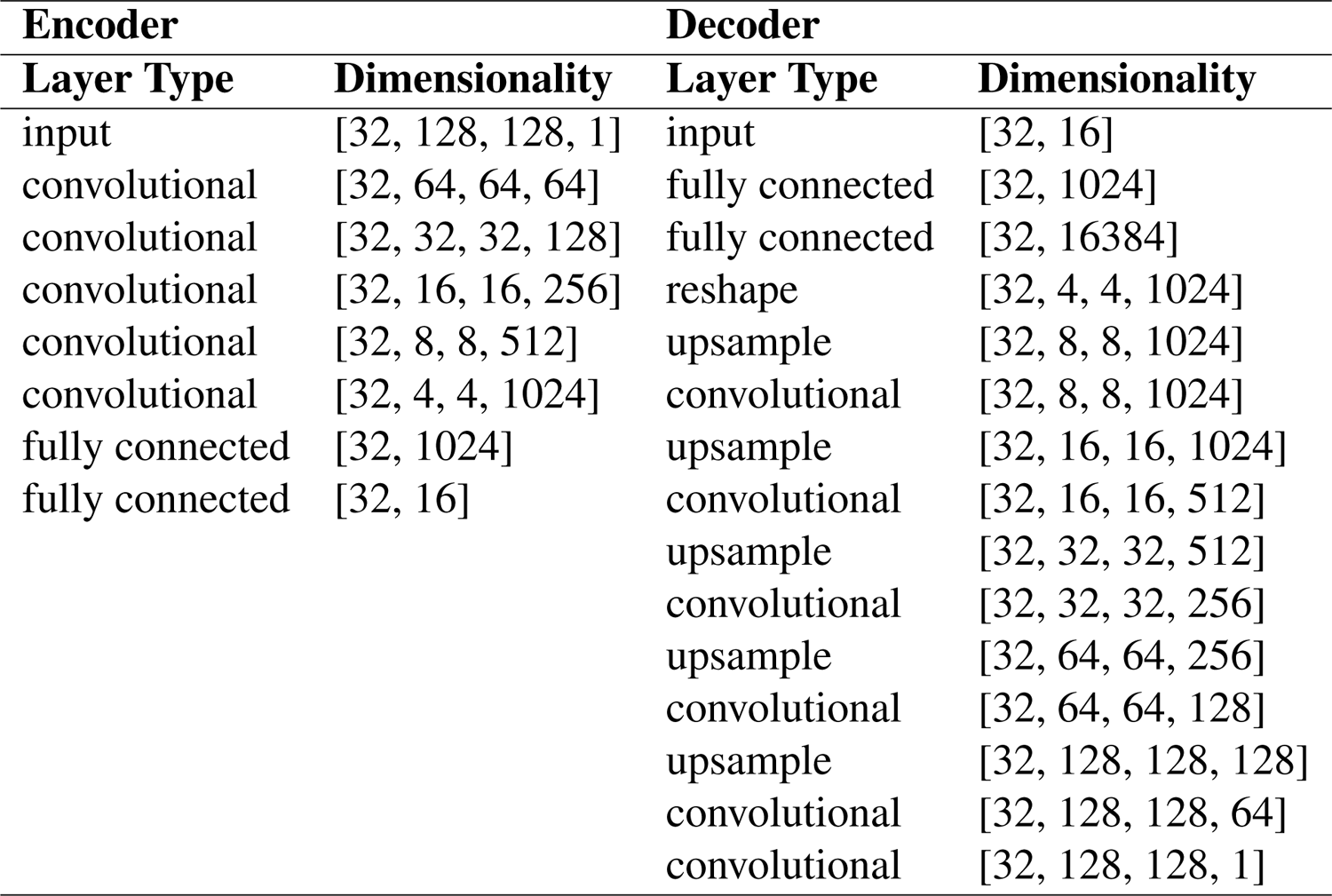
Variational autoencoder architecture outline

| **Encoder** |  | **Decoder** |  |
| --- | --- | --- | --- |
| **Layer Type** | **Dimensionality** | **Layer Type** | **Dimensionality** |
| input | [32, 128, 128, 1] | input | [32, 16] |
| convolutional | [32, 64, 64, 64] | fully connected | [32, 1024] |
| convolutional | [32, 32, 32, 128] | fully connected | [32, 16384] |
| convolutional | [32, 16, 16, 256] | reshape | [32, 4, 4, 1024] |
| convolutional | [32, 8, 8, 512] | upsample | [32, 8, 8, 1024] |
| convolutional | [32, 4, 4, 1024] | convolutional | [32, 8, 8, 1024] |
| fully connected | [32, 1024] | upsample | [32, 16, 16, 1024] |
| fully connected | [32, 16] | convolutional | [32, 16, 16, 512] |
|  |  | upsample | [32, 32, 32, 512] |
|  |  | convolutional | [32, 32, 32, 256] |
|  |  | upsample | [32, 64, 64, 256] |
|  |  | convolutional | [32, 64, 64, 128] |
|  |  | upsample | [32, 128, 128, 128] |
|  |  | convolutional | [32, 128, 128, 64] |
|  |  | convolutional | [32, 128, 128, 1] |

We modeled subjects’ psychophysical behavior using a Bayesian integration framework (Fig 1F-I), treating the cue syllable as a prior probability over the morph continuum (Fig 1F) and representing the uncertainty over the sensory stimulus as a Gaussian probability distribution centered around the true location of the target syllable in the continuum (Fig 1G).

### Context-dependent shift in perceptual decision making

We fit a psychometric 4-parameter logistic function (Fig. 2A) to each subject’s classification behavior for each morph continuum. We used the parameters of the fit psychometric model to test two, non-mutually exclusive, hypotheses of how the cue may affect behavior. Under the Bayesian integration hypothesis, (Fig 1F-H) syllable classification will be modulated by integrating the likelihood imposed by the target syllable with the prior imposed by the sequential cue syllable, leading to a shift in the decision boundary (Fig 2A, inflection point) in the direction predicted by the cue (Fig 2B top). This would support a shift in categorical perceptual decision-making through the integration of the sensory signal and its sequential context. We also examined whether information from the cue and target syllables are treated independently, as evidenced by an overall shift in the probability of a left or right response, but not a shift in the decision boundary (Fig 2B bottom). Across each syllable continuum and for each bird, we observe robust shifts in the decision boundary (inflection point, Fig. 2D; t(350)=18.5, p=6.9e-54), indicative of Bayesian integration underlying context-dependent categorical perception. Also consistent with the Bayesian model, the magnitude of the inflection point shift increases as the cue’s prediction probability increases (Fig. 2E; r^2^=0.447, p = 1e-52, n=1050). To examine this shift more closely, we fit the Bayesian model to each bird’s behavioral data (for each continuum) and predicted the inflection point shift given each cue probability. The red dashed line in Fig. 2E depicts a linear regression showing the close correspondence between the observed shift in inflection point and that predicted by the Bayesian model. We also observed an overall shift in decision probability (Fig. 2C), suggesting additional independent, non-Bayesian effects of the cue.

**Figure 2:**
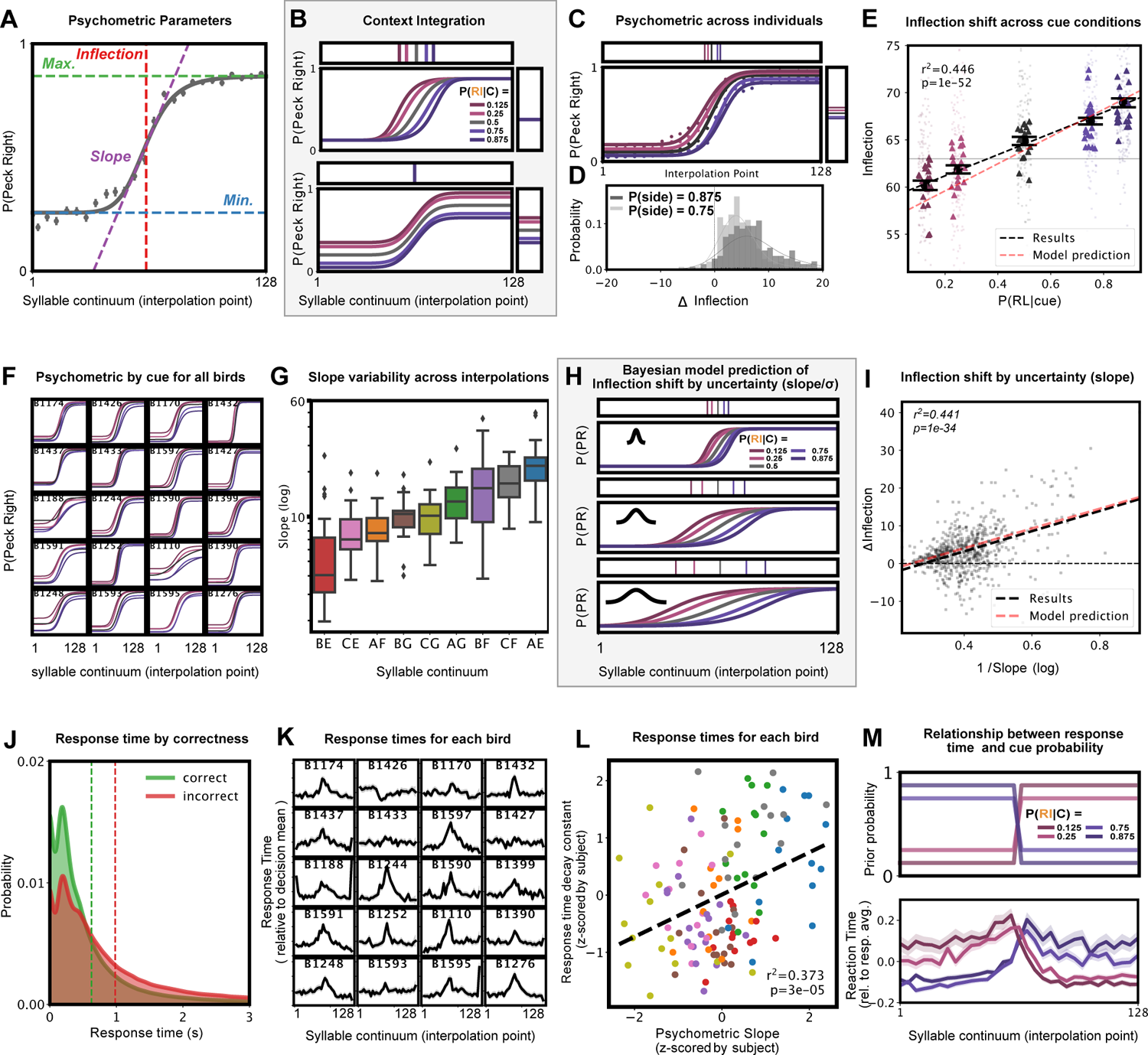
Behavioral support for the Bayesian model. (A) An example psychometric fit with parameters. (B; top) An example of the context-dependent category shift as a function of cue information hypothesis, as predicted by the Bayesian model. (B; bottom) An example of an alternative hypothesis, in which decisions are made either using the cue or the categorical stimuli, without integration of the two sources of information results in no category boundary shift. The corresponding lines in the connected horizontal and vertical boxes indicate the shift in the inflection point (vertical lines) as well as the midpoint between mid and max in the psychometric function (horizontal lines). Colors indicate the cue probabilities given in Figure 1. (C) The results across birds and morph indicate that both strategies from (B) are present in behavior. (D) Cue shift between left and right cues for each morph and bird at p=0.875 and 0.75. (E) The categorical boundary (inflection point) shifts as a function of the strength of the cue. The Bayesian model, predicts a similar shift from the uncued data. (F) Psychometric fits for cued conditions for each of the subjects. (G) Morphs (interpolations) vary on the slope of the fit psychometric function, indicating variation in uncertainty in decision making by morph. (H) The Bayesian model predicts a greater shift in categorical boundary as a function of the uncertainty of the categorical stimulus (*σ* of the likelihood and slope of the psychometric model). (I) As predicted by the Bayesian model, the shift in the categorical boundary increases as a function of uncertainty. (J) Response time across birds for correct versus incorrect trials. (K) Response time over the morph for each bird. (L) Decay constants of exponential decay fit to reaction time as a function of distance from decision boundary, in relation to the slope of the fit psychometric function, for each bird and morph. Point colors reflect the morph categories shown in panel (G). (M; top) The imposed prior probability in the task for each condition. (M; bottom) Reaction time over morph for each cue condition.

### Context-dependent perceptual shift increases with uncertainty

We observed substantial variation in the slope of the psychometric functions fit to each bird’s behavior. Some individuals drew a much sharper categorical boundary than others (e.g. B1432 vs B1110 in Fig. 2F) and the mean slope (averaged across individuals) also varies between syllable continua (Fig. 2G). The slope of the psychometric reflects uncertainty in the Bayesian model. Under greater uncertainty about the target syllable, the Bayesian model predicts that integration with the cue stimulus will result in a greater shift in categorical perception (i.e. the inflection point; Fig. 2H (*21*)). Consistent with this, we observed a smaller inflection point shift in the direction of the cue as the slope of the psychometric steepens (Fig. 2I, r^2^=0.441, p=1e-34, n=700), which again matches the quantitative prediction of the model (Fig. 2I, red dashed line).

### Reaction time represents likelihood and prior probability

Although not directly captured in the Bayesian decision model, we also expect that response times should reflect both the uncertainty in decision making (the slope of the psychometric) and the prior probability given by the cue syllable. Consistent with this, response times were longer on incorrect trials than correct trials (Fig. 2J; *t*(1979805) = 224.4, p *<* 1e-5). Likewise, response times for most (17 of 20) subjects increased on trials with target syllables closer to the categorical boundary, suggesting increased difficulty in classification (Fig. 2J). For each bird and syllable continuum, we fit an exponential decay model of reaction time as a function of distance from the categorical boundary. For syllable continua where a decay was observed (set at an r^2^ *>* 0.001 and decay range *>* 0.1 standard deviations), we found a strong relationship between the exponential decay constant, and the psychometric slope (Fig. 2L; *r*^2^ = 0.373, p=2e-5, n=129). Finally, we observe that response times are proportional to the prior probability imposed by the cue. Across subjects, response times are fastest when the target syllable is preceded by the strongest valid cue, and slowest when preceded by the strongest invalid cue (Fig. 2M; *r*^2^=-0.07, p*<*1e-5, n=1979805).

### 2.2 Physiological results

We recorded extracellular neural spiking activity using 1-2 (unilaterally or bilaterally) implanted 32-64 channel 1-8 shank silicon electrode arrays from freely behaving subjects (N=10) while they completed trials on the syllable categorization task and passively listened to the same stimuli. We targeted electrode arrays to the secondary auditory forebrain regions CM (Caudal Mesopallium), NCM (Caudomedial Nidopallium), and NCL (Caudolateral Nidopallium), and the primary auditory region Field L (supp. Fig 9. We recorded from a total of 13,854 putative single-neurons (See 4.17) distributed across the four brain regions (supp Fig 11). We clustered the waveform templates of analyzed units into three sub-types (supp Fig 10), characterized predominantly by their spike width (supp Fig 10).

### Quantifying response similarity and estimating a neurometric

We analyzed spike train data as spike vectors over the different syllable continua, by convolving the time histogram (bin width=10ms) of the stimulus-aligned spike train for each trial with a Gaussian kernel (*σ*=25ms; Fig 13). Figure 3E and F show sample spike trains and trial-averaged spike vectors for a sample unit for each syllable continuum. From the trial-averaged spike vectors, we computed a cosine similarity matrix between spike vectors for each syllable on each continuum (Fig 3I) from which we then computed a neurometric function (Methods, Fig 3J). We also used the cosine similarity matrix to compute a metric for each unit’s categorical responsiveness (Fig 3K-L; see Methods 4.23) reflecting the similarity of the unit responses within versus between syllable categories. From the original 13,854 auditory units we identified 7,994 units with categorically-relevant responses to the syllable continua (See Methods 4.24). On average, the spike vector responses for these categorical units varied smoothly across the syllable continua, but the degree of this smoothness varied (Fig 3M-N).

**Figure 3:**
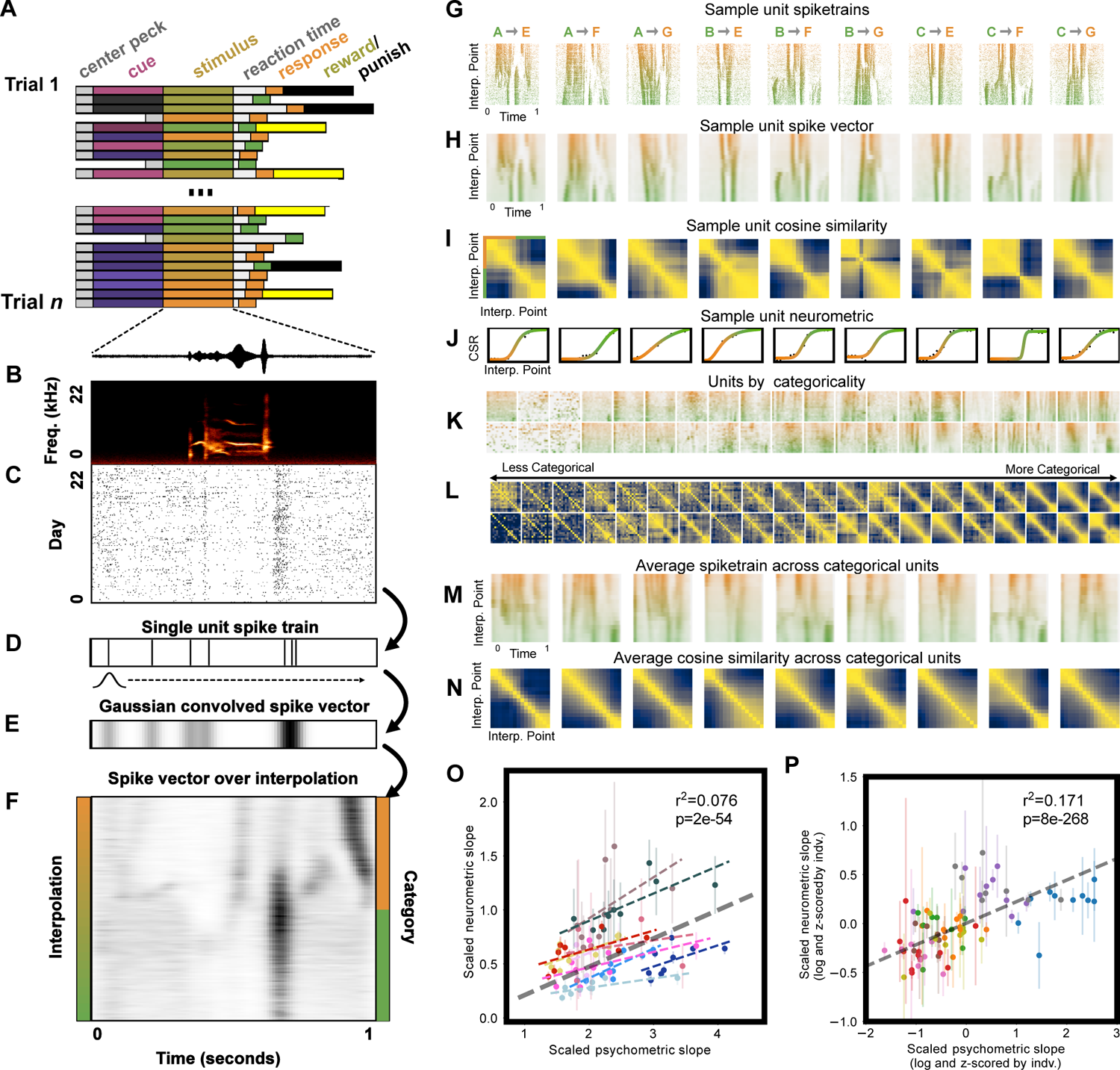
Neurometric functions of single units reflect psychometric functions of perceptual behavior. (A) Trial-by-trial behavioral data structure. (B) Spectrogram of categorical (morph) stimulus for a single trial. (C) Spike raster for a single unit across trials. (D) A single example spike train from (C). (E) A spike vector is computed as the spike train from (D) convolved with a Gaussian kernel. (F) The average spike vector for the unit in (C-D) for a single morph (A → E). (G) Sample spike trains for one unit across 9 morphs. (H) Spike vector representations of the spike trains from (G). (I) Cosine similarity matrices computed from the spike trains in H. (J) Neurometric functions are computed from the similarity matrices in (I). CSR stands for Categorical Similarity Ratio (see Methods 4.22). (K) Sample morph spike vectors (as in (H) for units, sorted by unit categoricality. (L) Similarity matrices for the units in (K). (M) Average spike trains across each categorical unit for morphs. (N) Average cosine similarity matrices across all categorical units, for each morph. (O) Psychometric slope (log transformed and scaled by psychometric range) versus neurometric slope (log transformed and scaled by psychometric range) for each subject and morph. Each subject is shown with a unique color and regression line. Regression line for pooled data is shown in gray. (P) The same data as in (O) z-scored by subject, where color corresponds to syllable continua (as in Fig 2).

### Neurometric slope reflects uncertainty

We compared the slope of the neurometric function to the slope of the psychometric function for each bird and syllable continuum (Methods). Across birds we observed a significant positive correlation between the slopes of the neurometric and psychometric functions (r^2^=0.076, p=2*e −* 54, n=41181), Fig 3P), which was consistent in 9 of the 10 birds. Controlling for individual variability by z-scoring the psychometric and neurometric slopes by bird strengthens this relationship (r^2^=0.171, p=8*e −* 268, n=41181), and reveals that across birds the same syllable continua occupy similar relative neurometric and psychometric positions (Fig 3Q). To assess the relationship between the psychometric and neurometric slopes directly, we performed a hierarchical regression comparing a prediction of the neurometric slope from the syllable continuum alone (`neurometric slope` *∼* `morph + subject`) to a prediction of the neurometric slope from the continuum and the psychometric slope (`neurometric slope` *∼* `morph + psychometric slope + subject`). We find that the psychometric function explains significantly more variance in the neurometric function than stimulus alone (*F* (1, 41163) = 21.9, *p*=3e-06, ΔAIC=19.9), suggesting that neural responses reflect behavioral variability in categorical perception across subjects and continua. Recall that the slope of the psychometric function is modulated by the likelihood, i.e. stimulus uncertainty, in the Bayesian decision-making model. It follows, therefore, that these categorical neural responses can be interpreted as carrying information about the stimulus uncertainty.

### Context modulates neural response

We have shown behaviorally that perceptual classification of song syllables can be biased by sequential context. As an initial assessment of whether cue syllables modulate neural responses to target syllables, we measured the overall spike rate change in each unit as a function of the predictive cue syllable in active behavioral trials (Fig 4). Controlling for spike rate variability across units, we find a strong effect of the cue syllables. Namely, the presence of a cue syllable significantly suppresses the z-scored spike rate evoked by the target syllable (*X*^2^(4, N = 857301) = 15196, *p <*1e-5; Fig 4A). This suppression is consistent across the motif continuum, stronger in active trials than passive playbacks (t(296464) = 46.5, p *<* 1e-5; Fig 4B), and is most prominent early and continues throughout much of the target stimulus playback (Fig 4C). Moreover, the magnitude of the cue-depended suppression is consistent within cue condition, and persists throughout stimulus playback, while in passive playback trials any cue-dependent effects quickly diminish (Fig 4D).

**Figure 4:**
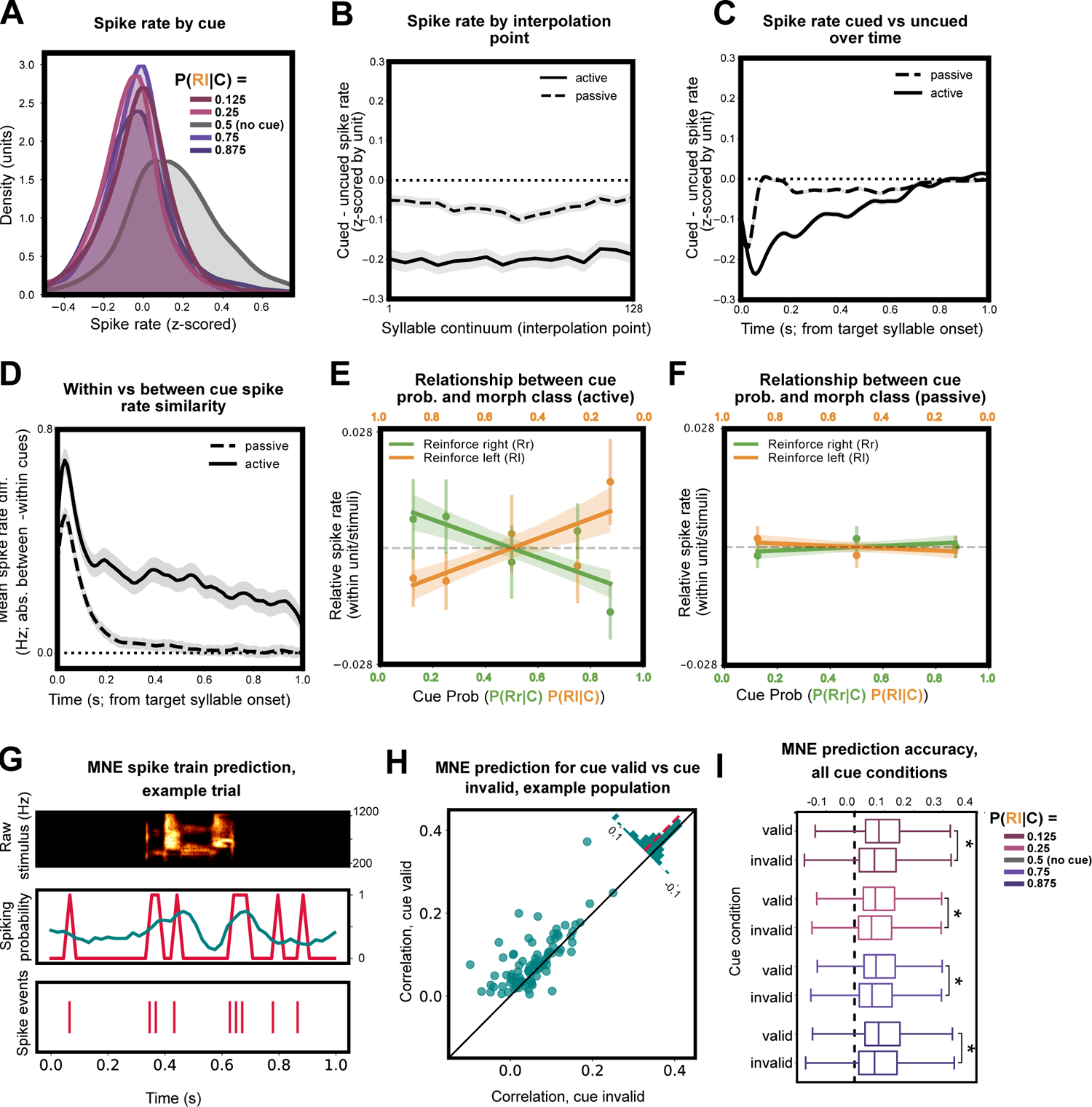
Predictive syllables modulate response to target syllable. (A) Differences in spike rate for each predictive cue syllable for active behavior playbacks, where uncued trials yield the highest spike rate. Spike rate is z-scored within each unit, and the difference is shown as the difference within stimulus across cueing conditions for each unit. (B) Spike rate suppression across units for cued trials by interpolation point. (C) Spike rate suppression across units for cued trials by time. (D) Within-cue vs between cue spike rate similarity over time. Mean difference in spike rate between cue conditions minus mean difference within cue conditions across time for passive and active trials. A value above zero indicates greater similarity within cues than between cues. (E) Relationship between cue probability and morph class, within unit and stimulus, measured through spike rate across morph playback. For both the left and right morph stimuli, a regression line is shown with a 95% bootstrapped confidence interval. (F) Same as (E) for passive trials. (G) A sample MNE receptive field prediction. (top) Raw spectrogram of the target syllable on an individual trial. (middle) Actual (red) and receptive field model predicted (teal) spiking probability (same trial). (bottom) Raster plot of spiking events (same trial). (H) Correlation values between actual and predicted spiking for cue valid vs. cue invalid trials. Trial correlation values were averaged across valid or invalid trials for each unit on an example recording day (N = 98 units). (I) Box-plots for distribution of trial averaged correlation values (as in H) for all units broken down by cue validity and strength. (* indicates significantly increased correlation value for valid verses invalid trials, post hoc t-test, Cue 0.125, *t*(9078) = 19.5*, p <* 0.001; Cue 0.25, *t*(9377) = 18.2*, p <* 0.001; Cue 0.75, *t*(9379) = 18.6*, p <* 0.001; Cue 0.875, *t*(9101) = 17.0*, p <* 0.001).

### Expectation suppresses spike rate in predicted stimuli

Because cue syllables are differentially informative (i.e. establish different priors) for upcoming target syllables, we reasoned that the magnitude of cue-specific response suppression might co-vary with the strength of this predictive information. To quantify this, we measured the interaction between cue informativeness and the category of the target syllable, while controlling for differences in each unit’s response between syllable continua (Fig 15). That is, how is each unit’s response within a syllable continuum modulated by the strength of the predictive cue? During active behavior trials, we find a significant interaction between cue strength and the syllable category. Across our sample, as the predictive strength of the cue syllable increases, spike rate decreases (*X*^2^(1, N = 857301) = 399, *p <*1e-5; Fig. 4E). This effect is abolished during passive playback (Fig. 4F) and in shuffle controls (Fig S15). We also examined the interaction between cue strength and syllable category in each brain region, neuronal sub-type, across individual syllable continua, and in each subject (Fig. 16). The effects were broadly consistent across our three unit sub-types and across the individual syllable continua (Fig. 16). Among the different forebrain regions represented in our datasets, Field L, NCL, and CMM all show similar effects, although the effect in CMM is weak. In NCM, we observe the opposite interaction: increasing cue strength facilitates target syllable responses (Fig. 16D). The effects in individual subjects mirrored the brain regions targeted in each (Fig. 16C).

### Expectation modulates encoding

We next assessed how cue-dependent response modulation was related to target syllable encoding in single neurons. For all categorical neurons in our sample, we fit a receptive field model (*22*) for each cue condition using a subset of trials across all continua on which the target syllable category matched the prediction of the cue (cue-valid trials). We hypothesized that cues modulate receptive fields to improve the encoding of likely upcoming syllables. If true, our receptive field models should provide a better prediction of the neural responses to the same target syllable on cue-valid compared to cue-invalid trials (i.e., trials where the target syllable category did not match the cue prediction). To test this hypothesis, we compared correlations between predicted and actual spike trains for the same target syllables on an equal number of held out cue-valid and cue-invalid trials (Fig 4G-H). Across all cue strengths, the receptive field models provided significantly better (more accurate) predictions of responses to target syllables on cue-valid trials than on cue-invalid trials (Linear Mixed Effects, cue validity: *β* = 0.015, *SE <* 0.001, *z*(73878) = 41.074, *p <* 0.001; Fig 4G-I). Although we cannot determine if encoding is enhanced by valid cues and/or degraded by invalid cues, the relative differences between these conditions supports the idea that receptive fields are rapidly reorganized by contextual cues in anticipation of likely upcoming syllables.

### Predictive response modulation is consistent with a shift in the likelihood of Bayesian model

Having established that neural responses to song syllables are modulated by predictive cues, we next explored how cue-related modulation reflects the categorical similarity between neural responses. We considered two hypotheses based on the Bayesian model presented in Figure 1F-I. The first follows a common characterization of categorical phoneme perception called the Perceptual Magnet Effect (*23*), whereby speech perception is warped around categorical boundaries to reduce discriminability of within category sounds. Within the Bayesian model, this perceptual warping results from the integration of prior distributional information with a noisy representation of the acoustic signal, yielding a shift in the posterior toward higher probability regions of acoustic space (*5*) (Fig 5A). In the context of our task, increasing predictive probability toward one side of the syllable continuum (i.e. in the context of a predictive cue) should lead to two outcomes: the within-category similarity of the posterior on the predicted side of the continuum should increase, and the within-category similarity of the low-probability side of the continuum should decrease (Fig 5B). Accordingly, if the neural responses in our samples reflect this posterior distribution their similarity should also increase as a function of predictive cue strength (Fig 5C,D). Changes to the posterior distributions in the foregoing model reflect perceptual decisions, but do not account for direct modulation of the stimulus likelihood. In visual systems, however, early sensory modulation in the context of predictive information appears to be qualitatively different than that in decision-making regions (*24*). As an alternative to the perceptual magnet effect model, therefore, we propose an extension to the Bayesian model in which predictive information directly modulates the stimulus likelihood (Fig 5). In this model, predictive information reorganizes sensory representations to increase acuity in regions of acoustic space where events are more likely to occur. We model this as a narrowing of the likelihood (here, *σ* of the Gaussian distribution) reflecting the redistribution of neural resources to reduce perceptual noise in predicted regions of stimulus space. This model makes the opposite predictions to those focused on the posterior distribution (Fig 5F). That is, increasing acuity (i.e. decreasing noise in the likelihood) in high-probability regions of stimulus space should *reduce* within-category similarity (Fig 5G,H). Accordingly, if neural signals in our samples reflect the stimulus likelihood, their similarity should decrease as a function of predictive cue strength.

**Figure 5:**
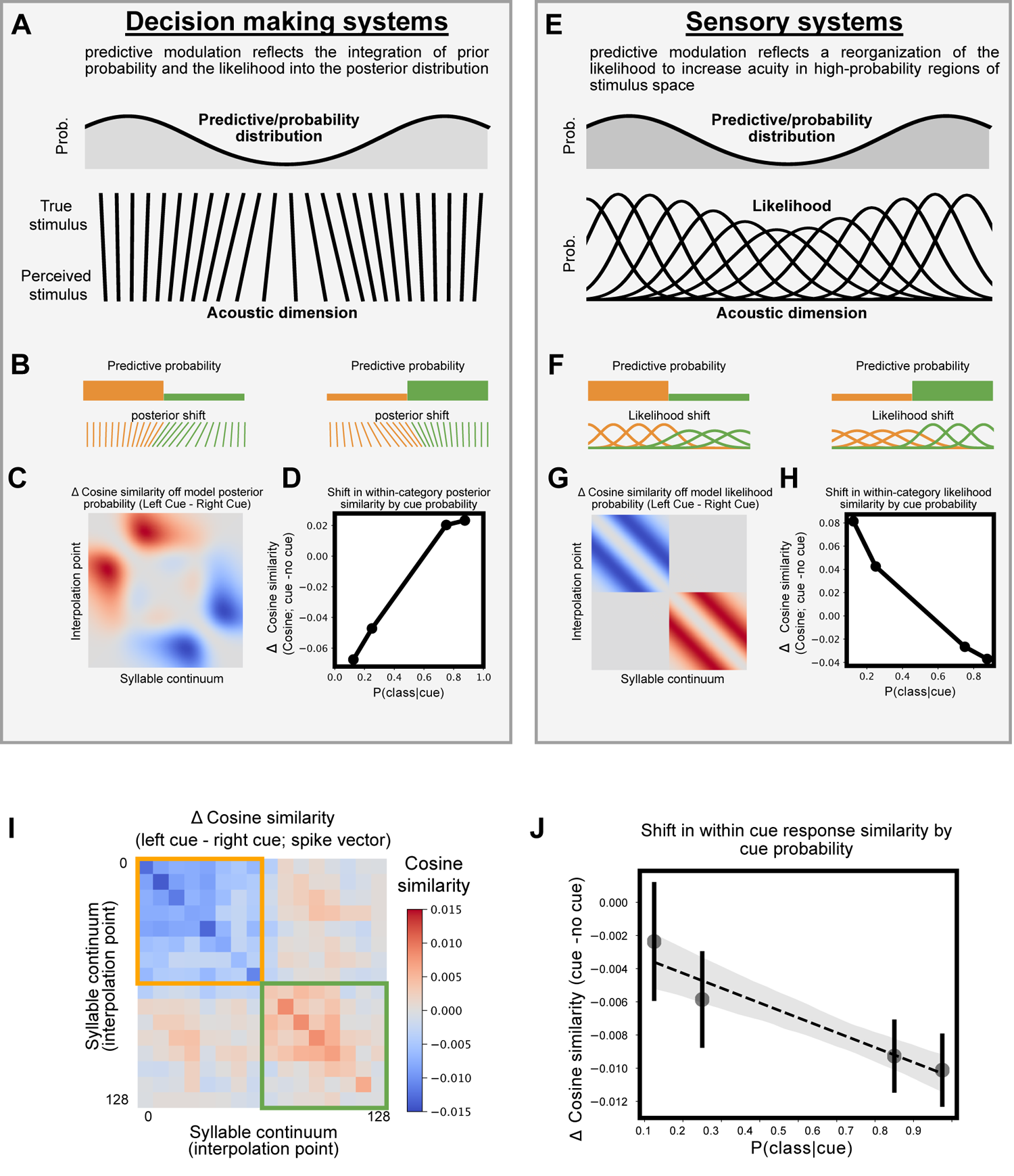
Modulation of response similarity as a function of predictive cue probability. (A) A summary of the Perceptual Magnet Effect (*23*) and the corresponding Bayesian model (*5*). (B) Visualizing the posterior shift from (A) in the context of this manuscript’s behavioral experiment. (C) The shift in the Bayesian model’s posterior distribution (measures as cosine similarity over the PDF) when cued. (D) The shift in the within-category posterior cosine similarity as a function of the probability of that class. (E) An extension to the model in (A) in which predictive information biases the likelihood to increase acuity in high probability regions of acoustic space. (F) Visualizing the likelihood shift from (E) in the context of this manuscript’s behavioral experiment. (G) The shift in the Bayesian model’s likelihood distribution (measures as cosine similarity over the PDF) when cued. (H) The shift in the within-category likelihood cosine similarity as a function of the probability of that class. (I) The observed shift in spike train vector cosine similarity for left-cued minus right-cued trials. The shift is depicted here averaged across units and morphs. (J) The relationship between the probability of the stimulus class and the shift in similarity from baseline (the uncued condition).

To test these hypotheses and assess whether the similarity of neural responses changed as a function of the cue, we compared the trial-to-trial cosine similarity of the spike vector response across syllable continua. Our results support the model of likelihood modulation rather than posterior integration. We find that in the presence of a predictive cue, the within-category similarity is higher in the non-predicted class than the predicted class (Fig 5I; t(1538963) = 12.8, p=2.07e-37). Moreover, the within-category similarity across units and continua decreases sig-nificantly as a function of the probability of the cue class (*r*^2^ = −0.020, *p*=1.06e-10; n=108153; Fig 5J). This pattern of results suggests that perceptual acuity is selectively sharpened over predicted regions of acoustic space, with the goal (we contend) of decreasing representational noise. As in our spike rate analyses, we repeated the similarity analysis over individual morphs, brain regions, subjects, and unit sub-types. Results were broadly consistent across syllable continua and unit sub-types (Figs 18A,B, 19A,B). Across brain regions, our results parallel those in the spike rate analyses (Figs 18, 19). We observe a decrease in similarity between predicted syllables in CMM, Field L, and NCL, while in NCM we do not observe a change in similarity (Figs 18D, 19D). In individual subjects, the effects mirrored the brain regions recorded (Figs 18C, 19C).

## 3 Discussion

Categorical perception is fundamental to a wide range of cognitive behaviors. It involves a non-linear mapping between graded physical sensory signals and behavioral decisions. In real-world conditions, this mapping is not fixed but can vary between groups and individuals, and within individuals across developmental and short time-scale temporal contexts in both human speech (*2*) and bird song (*9*). The neural mechanisms underlying categorical perception and its modulation are not well understood. We trained songbirds on a categorical perceptual decision-making task, controlling the predictive contextual information in sequences of vocal elements. We showed that songbirds use this information to bias categorical perception of their vocal signals. We then show that the qualitative and quantitative features of this bias are captured by a Bayesian model of perceptual decision-making, similar to that proposed for human context-dependent categorical speech perception (*4, 5*), in which the degree of bias reflects the integration of the predictive contextual information and uncertainty over natural stimulus dimensions. Using the same behavioral framework, we then recorded from populations of sensory neurons while birds were making categorical perceptual decisions. We found that many sensory neuronal responses throughout the auditory forebrain reflect the categorical structure of the natural stimulus space (Fig 3) and that syllable sequence predictability biases sensory representations by suppressing spike rates, modulating syllable encoding (Fig. 4), and reducing stimulus uncertainty (Fig. 5). We show that these neural responses are consistent with a Bayesian model for categorical perception in which predictive cues act to modulate stimulus likelihood.

Ongoing work in speech seeks to uncover the neural systems for processing predictive information relevant to lexical and pre-lexical feedback (*4*). In animals, prior work across sensory modalities and behavioral paradigms establish that predictive information increases responses in motor and decision-making related brain regions, while in sensory regions, the opposite effect is observed: activity tends to be suppressed when expectation is increased (i.e. expectation suppression; (*24–27*)). We suggest that this response suppression marks a rapid shift in sensory processing (*28*) that enhances the encoding of likely upcoming events, perhaps by highlighting specific stimulus dimensions or otherwise reducing internal representation noise. This rapid sensory modulation is likely to have multiple downstream effects, including as we show here, the categorical perception of natural communication signals in vocal sequences.

## Acknowledgments

This work was supported by a CARTA Fellowship to T.S., NIH 5T32MH020002-20 to T.S., and 5R01DC018055-02 to T.G.

## 3.1 Data and code availability

Data will be available upon publication. Code is available at https://github.com/ `timsainb/cdcp_paper`.

## 3.2 Author contributions

TS and TQG designed the experiments. TS, TM, EA, SR, MT, BT, PTM, and MT carried out the experiments. TM performed the MNE analyses and TS performed all other analyses. TS, TM, and TQG wrote the first draft of the paper. All authors made contributions to the final version of the paper and interpretation of data.

## 4 Methods

### 4.1 Summary

Experiments consisted of a behavioral component and a chronic physiology component. The experimental protocol for the behavioral component was kept constant by using the same software and hardware in both conditions, with the addition of chronic electrophysiological recording in the physiology component.

### 4.2 Subjects

Behavioral data were collected from 20 wild-caught European starlings of unknown sex. Before beginning experimental training, subjects were housed in a large mixed-sex aviary. Of the 20 starlings used for behavior, 10 individuals were used for chronic physiology.

### 4.3 Ethical note

All procedures were approved by the Institutional Animal Care and Use Committee of the University of California (S05383).

### 4.4 Datasets

Our final behavioral dataset was composed of 4.8 million behavioral trials from 20 birds. Our final chronic neural dataset was composed of 402,797 behavioral trials, with 365,360 responses, a total of 1,594,257 audio playbacks, occurring over 5,345 hours (222 days) of recording, across 10 birds.

### 4.5 Stimulus generation

Stimuli were syllables of European starling song synthesized from a Variational Autoencoder (VAE) trained on syllables extracted from a library of European starling song (*29*).

#### Training dataset

Syllables were segmented from the full songs of starlings with the dynamic thresholding approach outlined in (*30*) and available in the vocalization segmentation python package (https://github.com/timsainb/vocalization-segmentation). Syllables were zero-padded symmetrically at their beginning and end to be 1 second long. Spectrograms of each syllable were computed with 128 frequency bands spaced between 50 and 22050 Hz, and downsampled to 128 time-bins (128 Hz), resulting in a 128×128 spectrogram of each syllable, used to train the VAE.

#### Neural network

The neural network architecture we used followed those in our AVGN repository (https://github.com/timsainb/avgn_paper). We used a convolutional VAE architecture with a 16-dimensional latent space. The network was trained on batches of 32 syllables at a time. Artificial neurons used a leaky ReLu non-linearity. The network was trained with the ADAM optimizer in Tensorflow.

#### Sampling and synthesis

Each syllable stimulus (used for cues and endpoints) was sampled from the original dataset and passed through the VAE. The stimuli were chosen to be diverse, well-reconstructed in the VAE, and roughly equidistant both in spectrogram space (both input and reconstruction) as well as the latent space of the VAE. It is not expected that distances in spectral or neural network latent space would have a 1:1 relationship with an animal’s perception of similarity. Morph syllables were sampled from 126 evenly spaced points only the linear interpolation between the latent (16D) representations of a pair of endpoint syllables and passed through the decoder of the VAE, producing the final 128 syllable continuum including the two endpoint syllables. Waveform stimuli were then generated from the spectrogram output of the decoder of the VAE using the Griffin-Lim algorithm. These waveforms were the stimuli used for playback to the birds.

### 4.6 Behavioral training paradigm

Birds were initially trained to differentiate between the two syllable endpoints for a single continuum. After several days of above-chance accuracy with one pair of syllable endpoints, the number of endpoints was increased until the birds showed above accuracy classification of the endpoints of all 9 continua. After learning the correct response for all endpoints, birds were transferred to the full stimulus set which included all 128 syllables (linearly sampled and equally spaced in latent space) spanning each of the 9 continua (1152 syllables total). After the birds were performing reliably above chance on each full syllable continuum for several days, we added cue syllables preceding the target syllables to provide context-dependent information at p=0.125, p=0.25, p=0.5, p=0.75, and p=0.875.

### 4.7 Training parameters

Several behavioral parameters were used in behavioral training, given here for reproducibility. Trials were reinforced on a variable ratio schedule between 2-4 responses, manually set for each bird to maximize the number of trials each day without the loss of more than 10 grams of weight from baseline when in the restricted feeding condition. Punishment was set at a 5-second lights-off period, during which new behavioral trials could not be initiated. A minimum of 1 second between trials, regardless of response, was imposed. Birds were given a 5-second window to respond after stimulus playback. Lighting conditions were set to match seasonal sunrise and sunset times in the experimental location (San Diego, California).

### 4.8 Cue stimulus

Like the morph syllables, the cue syllables are 1-second long, synthesized by reconstruction from the variational autoencoder. Behavioral trials were presented with one of 6 cue conditions: no cue P(L—No Cue)=0.5 (NC), cue with no predictive information P(L—Cue)=0.5 (CN), cue left at p=.875% P(L—Cue)=0.875 (CL1), cue left at p=0.75% P(L—Cue)=0.75 (CL0), cue right at p=.875% P(L—Cue)=0.125 (CR1), cue right at p=0.75% P(L—Cue)=0.25 (CR0). 16% of trials were presented in the no cue condition (NC). 4% of trials were presented with the uninformative cue condition (CN). The remaining 80% of trials were evenly split between the cue right and cue left conditions. Because the CN condition was sampled with a substantially lower probability than the other conditions, resulting in a low number of total trials in comparison to each other cue condition, it was not included in physiological analyses. In passive physiology playback conditions, due to time constraints in playing back the full stimulus set of 128 interpolation points for each of 9 morphs and 6 cue conditions, we played back only the 87.5% predictive cue conditions in the AE and BF morphs.

### 4.9 Psychometric fit

To assess the shift in categorical perception, in each of the birds (n=20) we fit a psychometric (four-parameter logistic) function both to the overall responses to stimuli in the left and right categories of the morph, as well as to each individual morph.

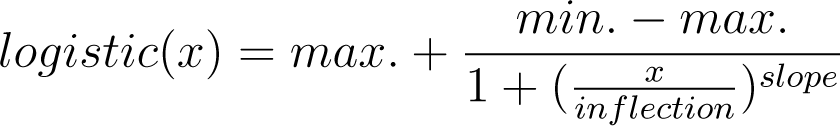

### 4.10 Bayesian integration hypothesis

To formalize our hypothesis, when a stimulus varies upon a single dimension *x*, the perceived value of *x* as a function of the true value of *x* and contextual *cue* information can be described by Bayes’ rule:

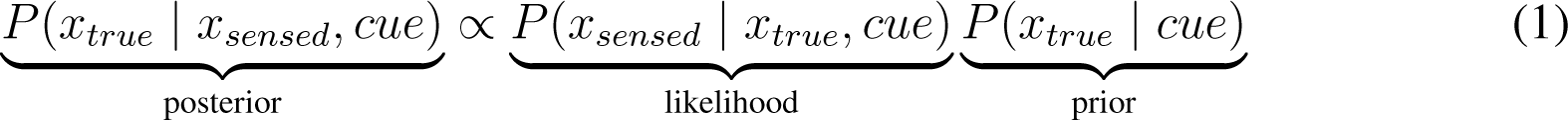

By modulating the prior distribution of the categorical stimuli (*x*) with a cue, we predict that the perception of *x* will shift.

Preceding each to-be-categorized target stimulus (*x*), the cue stimulus provides predictive information about the category of the target stimulus. By treating this cue stimulus as a prior probability over *x*, we predicted that the determined posterior probability of *x* given sensory information and the cue stimulus would shift the classification of stimuli near the boundary between the two classes in the direction predicted by the cue stimulus.

Explicitly, we treat the likelihood of a target being sensed *P* (*x_sensed_ | x_true_, cue*) as a Gaussian probability distribution around the true target *x_true_* (*5, 31*):

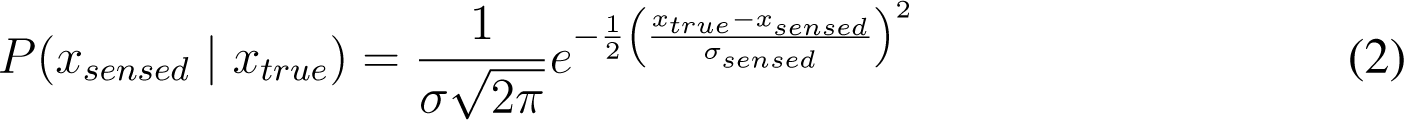

and shift the prior probability as a function of the cue

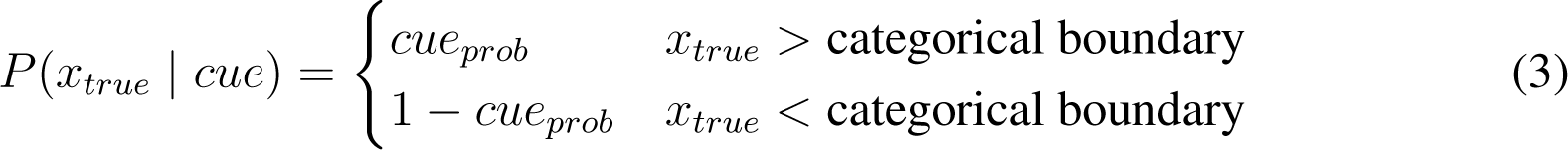

where *cue_prob_* represents the predictive probability of the cue stimulus. We predict that birds will make a categorical decision based upon the posterior,

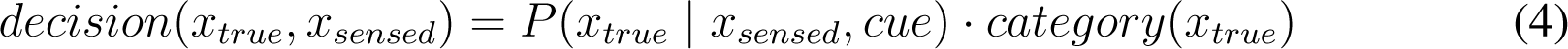

where *category*(*x_true_*) is simply the trained category label of *x* in the 2AC task:

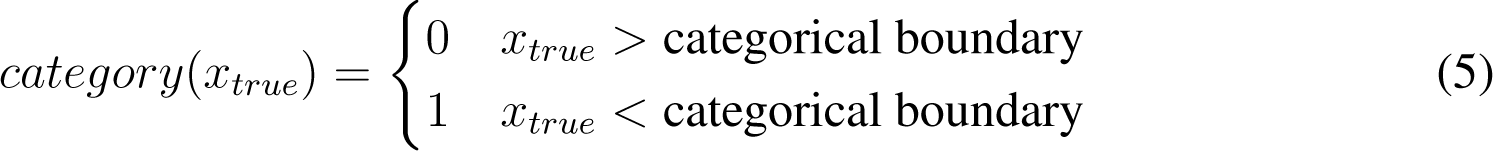

Under this model, the categorical decision of the bird is modulated by the prior cue information, resulting in a shift in the categorical decision point along the stimulus dimension in the direction predicted by the cue (Figure 1I).

#### Bayesian fit

In addition to fitting a psychometric function capturing the shape of the behavioral responses, we fit a Bayesian model reflecting our probabilistic hypothesis described above. This model used five parameters: the shape of the Gaussian of the likelihood (*σ_sensed_*), a parameter corresponding to side bias in the apparatus (*γ*), and parameters representing inattention to the cue stimulus (*δ*), the target stimulus (*β*), and overall inattention to the task (*α*).

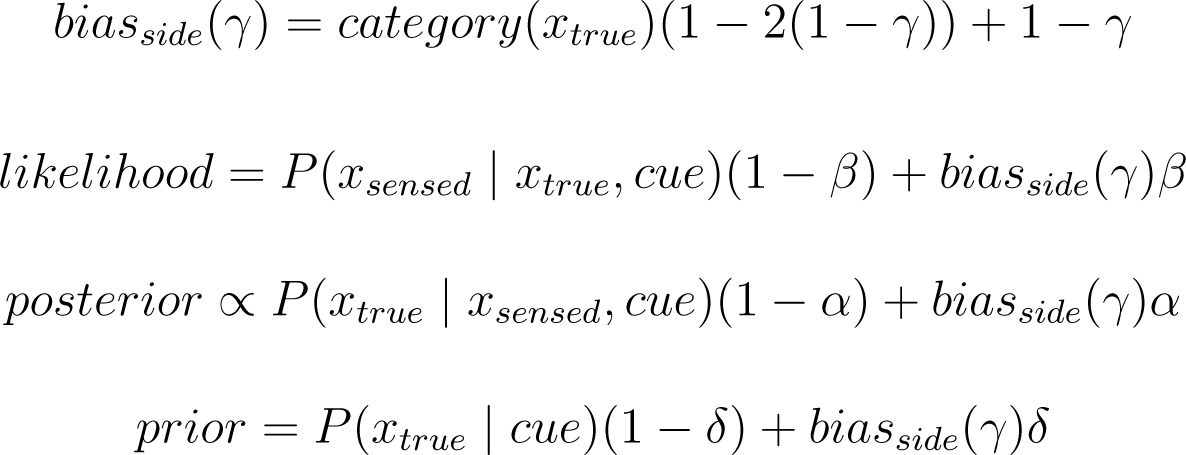

### 4.11 Response time

For each behavioral trial, we measured the time between the end of a stimulus presentation and the time that a subject’s beak was detected in a behavioral response port. In Figure 2J-M we found that the response time varied based upon stimuli and cue conditions. To account for side biases in decision making (e.g. the bird having a position preference when engaging with the behavioral apparatus that positions them further toward the left or right peckport, for each analysis we z-scored reaction time for each bird’s responses to each interpolation and class.

To parameterize the decay in response time as a function of the distance in the morph from the decision/class boundary, we fit the decay in response time to an exponential decay function (Fig 6). We discluded three birds from analysis (B1426, B1433, B1427) who we observed did not exhibit the same decay in reaction time as a function of distance from the decision boundary (Fig 2K).

**Figure 6:**
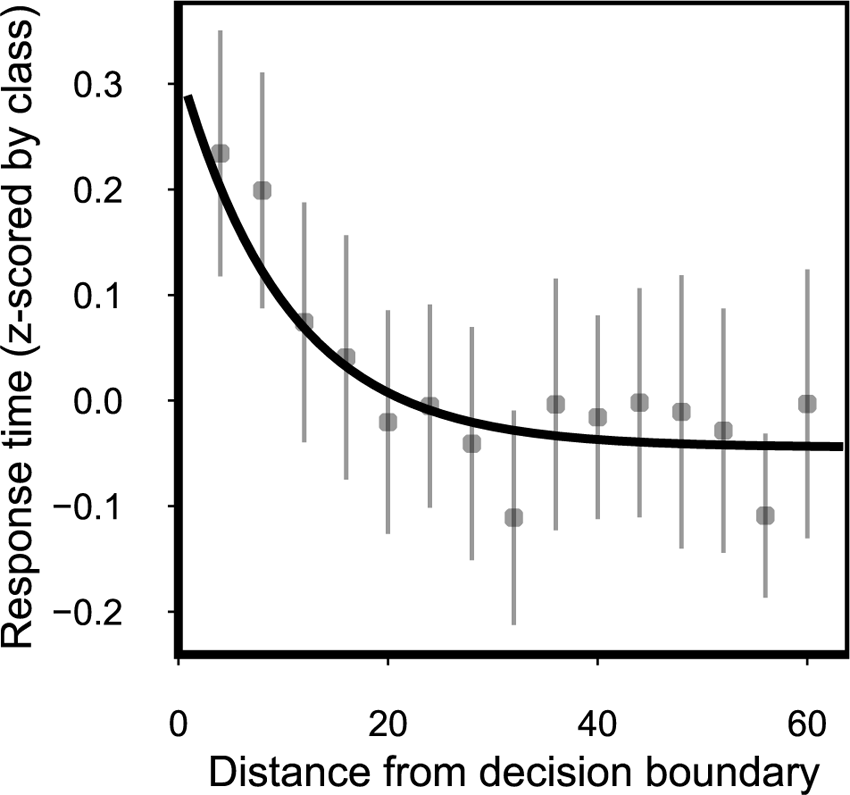
Sample fit of response time decay for a single morph (AE) for a single bird (B1174).

### 4.12 Chronic electrophysiology

We used 32 or 64-channel Neuronexus Si-Probes (A4×2-tet-7mm-150-200-121, Buzsaki32, Buzsaki64, A1×32-Edge-5mm-20-177) implanted either unilaterally or bilaterally. Probes were coated with PEDOT using an Intan RHD Electroplating Board no more than one week prior to implant. Probes were mounted on 3D-printed drives (described in Section 4.14), which were stereotactically implanted using the procedure outlined in Section 4.15. Extracellular voltages were amplified and digitized at 30kHz using an Intan RHD recording headstage, output through an SPI cable through an electrically assisted commutator to an Open Ephys recording system.

### 4.13 Behavioral neural acquisition interfacing with PiOperant

Behavioral and physiology were synced using a custom-designed Raspberry-Pi-based system (PiOperant) for automating our behavioral paradigm and interfacing with the OpenEphys neural acquisition device (Fig 7). PiOperant interfaces with our behavioral panel using the Python software pyoperant (https://github.com/gentnerlab/pyoperant). Behavioral states and audio signals were input and synced with OpenEphys over two HDMI inputs (digital and analog) and a ZMQ interface containing additional information about behavioral trials.

**Figure 7:**
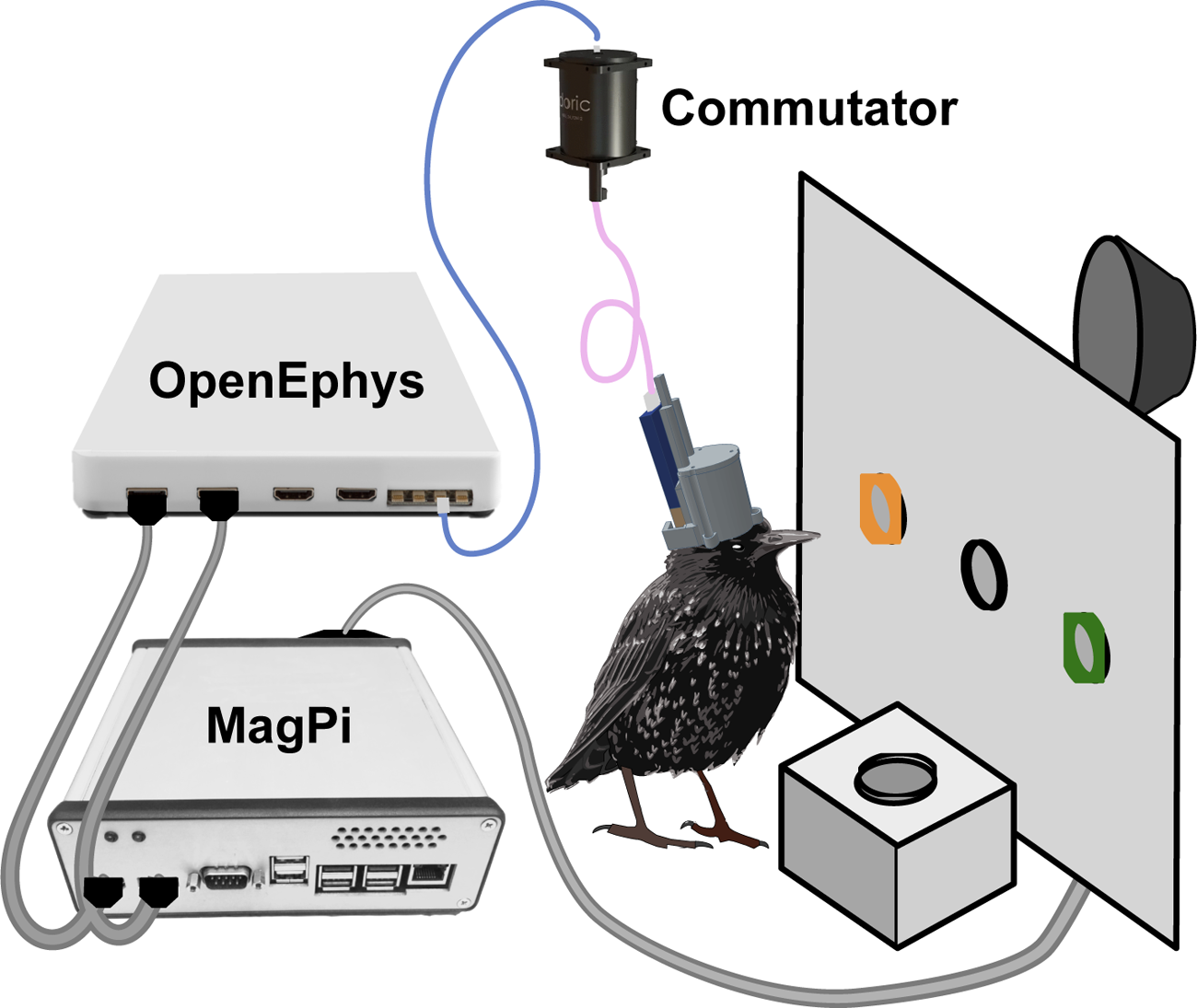
Continuous recordings are performed in freely-moving birds while recording physiology alongside operant conditioning behavior.

### 4.14 Microdrives and head caps

Microdrives and head caps (Fig 8) were custom-designed over the course of this experiment and were printed using a FormLabs Form3 3D printer using FormLabs standard grey resin printed at 25-50 micron resolution. Microdrives were comprised of a drive, a shuttle, and a MiniTaps 6/16” 00-90 gold screw, hand-tapped and fastened to the drive with a brass nut. The screw was used to raise and lower the shuttle manually, at a depth of 282 microns per full rotation. Head caps were designed to be removable and enable moving probes further down, as well as easy to explant allowing re-use of probes.

**Figure 8:**
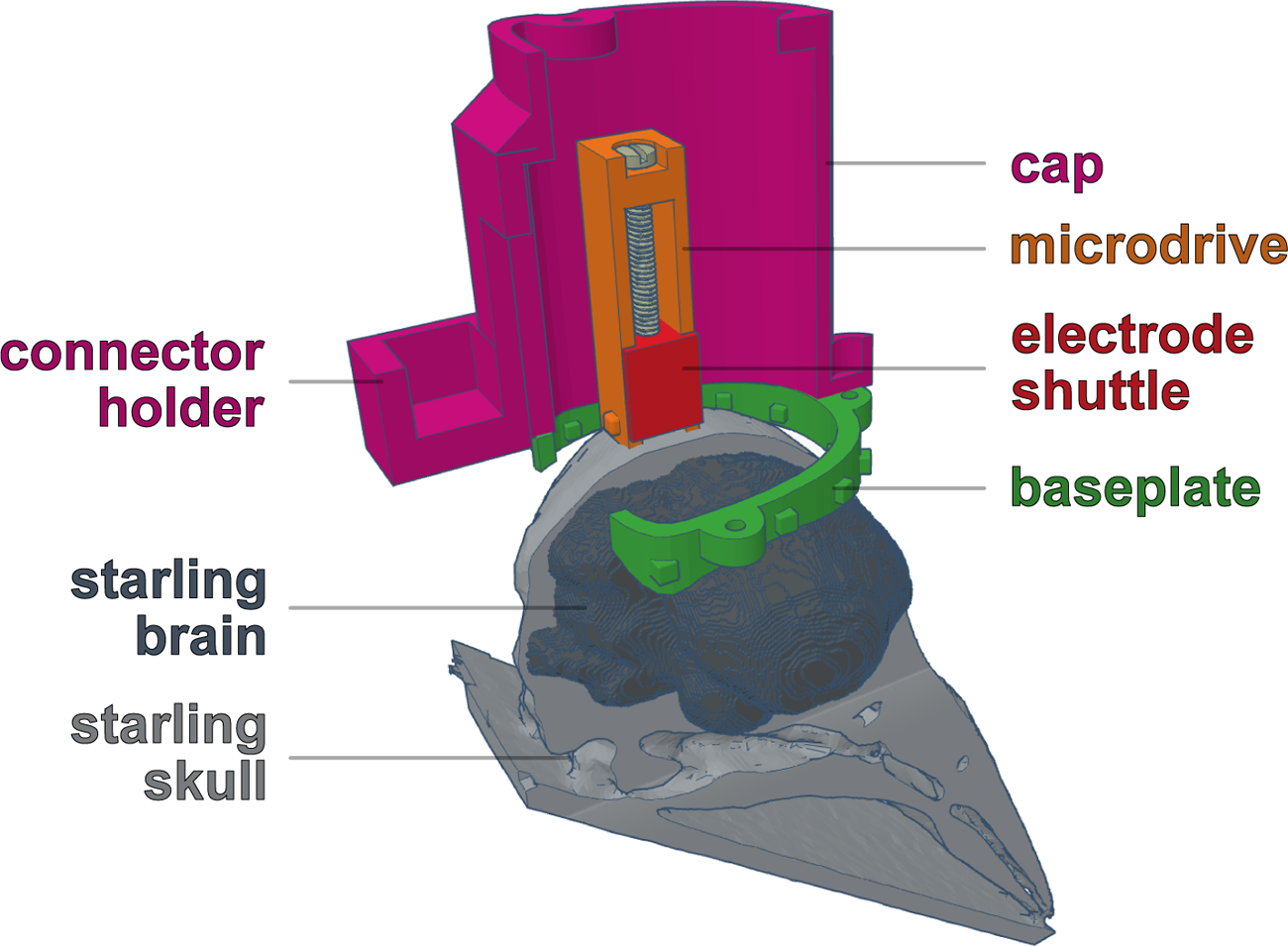
Headcap and microdrive design. Birds are unilaterally or bilaterally implanted with 32-64 channel electrodes using 3D printed microdrives and protective head caps.

### 4.15 Electrode implant procedure

Subjects were given analgesia by means of a 5mg/kg dose of carprofen (Rimadyl) administered intramuscularly. Animals were then anesthetized with a gaseous mixture of Isoflurane/oxygen (1-2.5%, 0.7 lpm). The scalp and feathers around the scalp were then removed and part of the skull over the y-sinus (the stereotactic reference sinus between the cerebellum and the two hemispheres of the brain) was visible. A craniotomy was opened above the recording site. A second craniotomy for the ground was then performed several millimeters away from the primary craniotomy. A platinum-iridium ground wire was then inserted in the craniotomy above the dura and glued to the skull. The baseplate for the head cap was then cemented (Metabond) to the skull. The durotomy was then performed in the original craniotomy, and the electrode was stereotactically lowered, attached to the microdrive at a rate of no more than 100 microns per minute. Once the final site was reached, the microdrive was then cemented to the skull, and a silicone base was applied above the craniotomy to prevent infection. The head cap was then screwed into the baseplate, protecting the recording site and probe. The headstage was then attached to the outside of the head cap.

In some individuals, multiple implants were performed in serial when one probe failed by explanting and removing the first probe and microdrive, creating a new craniotomy in the opposite hemisphere and durotomy, and implanting a new probe/microdrive. In one individual, two probes/drives were implanted simultaneously one in each hemisphere.

### 4.16 Recordings and behavior blocks

Recordings were performed 24 hours per day in order to track individual neurons over days. Recordings consisted of (1) behavior blocks, in which subjects freely interacted with the behavioral apparatus, (2) a free feeding period, in which the behavioral apparatus presented food to the bird without requiring the bird to perform trials, (3) a passive playback block, in which lights were turned off and the birds were passively presented with stimuli, and (4) a sleep block, in which the lights were left off and no stimuli were played back.

We recorded from 10 subjects over a total of 222 days (5317 hours) of recordings. Chronically implanted subjects performed over 400,000 behavioral trials during recording. In addition, during the evening after birds had completed their behavioral trials for the day we turned the lights out in the behavior boxes and passively played back the same morph stimuli to the birds totaling 1.2 million passive playbacks while recording.

#### Chronic behavior blocks

Chronic behavior blocks were matched to behavior blocks without physiology. The behavioral apparatus was left on throughout the day, allowing subjects to initiate trials through a peck in the central peck port. Trials were intermittently reinforced with a food reward and punished with the lights briefly turning off on incorrect trials. Using this paradigm, subjects performed several thousand trials per day.

#### Chronic passive playback blocks

At a set time at the end of each day, we turned the lights out in the bird’s operant conditioning block and passively played back the morph stimuli to the bird. The bird’s activity and sleep state during this time was not monitored. The silence interval between stimuli was randomly sampled between 1.1 and 1.5 seconds.

### 4.17 Spikesorting and merging over long-term chronic recordings

Spikesoring was performed over each 12 hour block of recording using Kilosort 2-2.5 (*32*) and SpikeInterface (*33*). LFP was bandpass filtered between 300 and 6000 Hz and further normalized using common median referencing. To retain units across days/sorts, we additionally used an overlapping procedure to merge each neighboring pair of recordings together. To do so, we took the last 30 minutes of the previous recording, and the first 30 minutes of the following recording, and separately sorted that hour-long recording, which overlapped with the two larger recordings. We then computed the overlap between units in the overlapping recording and each of the two full recordings. Units were then considered to be the same unit if their “agreement” score (SpikeInterface; the spike coincidence of the two units) was above a set threshold (set at 0.5). Units from each of the larger recordings that were merged with the same unit in the overlapped recording were then merged, allowing the same unit to be tracked over multiple days.

### 4.18 Stimulus alignment

Stimuli playback was aligned to neural data using a 1kHz sine way sent from the MagPi behavioral control device to the OpenEphys acquisition board collected simultaneously with neural data, alongside a binary switch indicating the onset and offset of playback. An additional message giving information about the specific trial was sent over the local network via ZMQ.

### 4.19 Localizing units

Unit locations were defined as the location of the peak recording channel on which the unit was present. The recording channel was determined from its position within the shank, and the shank’s position relative to the stereotactic implant. Stereotactic implant locations were recorded relative to the Y-sinus between the cerebellum and two hemispheres of the brain, and the depth relative to the surface of the brain. Implant locations relative to nuclei were then determined relative to voxel mapping of the European starling brain atlas (*34*), as shown in Fig 9A,B). Sample unit spike trains for each nucleus are shown in Fig 9C).

**Figure 9:**
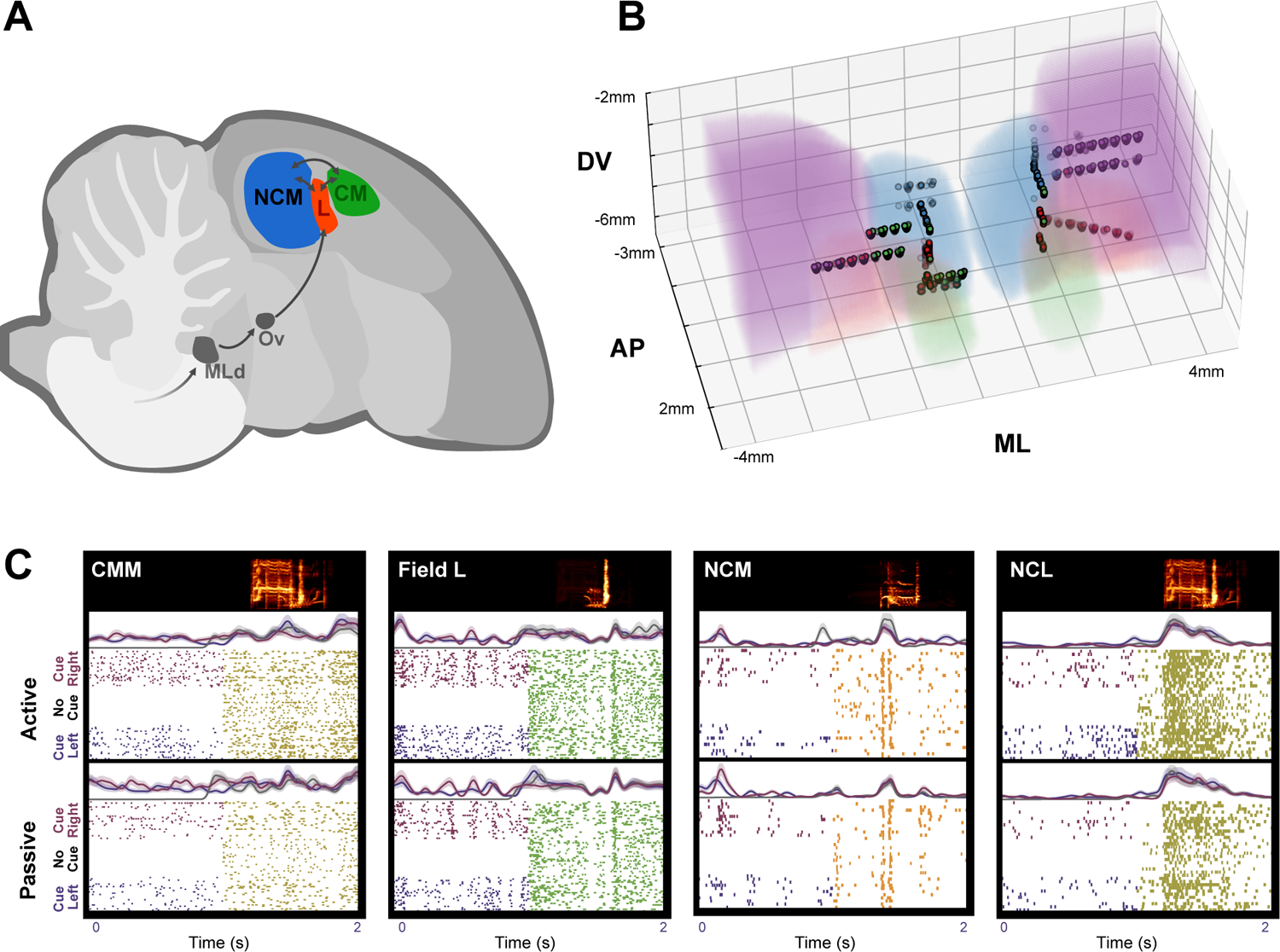
Recording sites. (A) Diagram of auditory input to the songbird brain. Nuclei OV projects to the primary auditory region Field L, which has bidirectionally projections with NCM and CMM. NCL (not pictured), lateral to NCM, additionally exhibits bilateral projections with Field L. (B) A visualization of recording sites, shown over top of the starling brain atlas (*34*). (C) The top of each panel shows a spectrogram of the morph stimulus played back. Below, a trace is shown for three cue conditions (No cue, P(Rl—C) = 0.125, and P(Rl—C) = 0.875) corresponding to the average Gaussian convolved spike vector and 95% CI for active trials. Below the trace are sample spike rasters for each cue condition, where each row is a trial. Below the rasters, the sample trace and raster plots are repeated for the same unit in the passive trial condition.

### 4.20 Clustering unit spike shapes

Unit spike shapes were clustered using the Birch clustering algorithm (*35*) fit to the template voltage trace of the peak channel of each unit into 3 clusters (Fig 10). The clusters found were primarily observed to vary upon their spike-width, and were (post hoc) manually labeled on the basis of spike width (i.e. wide, middle, narrow). Narrow spiking cells are generally believed to be inhibitory, while wide-spiking cells are thought to be excitatory (*36, 37*). Spike width was negatively correlated with spike rate (logged; r^2^ = −0.288, p=2.17e-129, n=7994; (Fig 10)).

**Figure 10:**
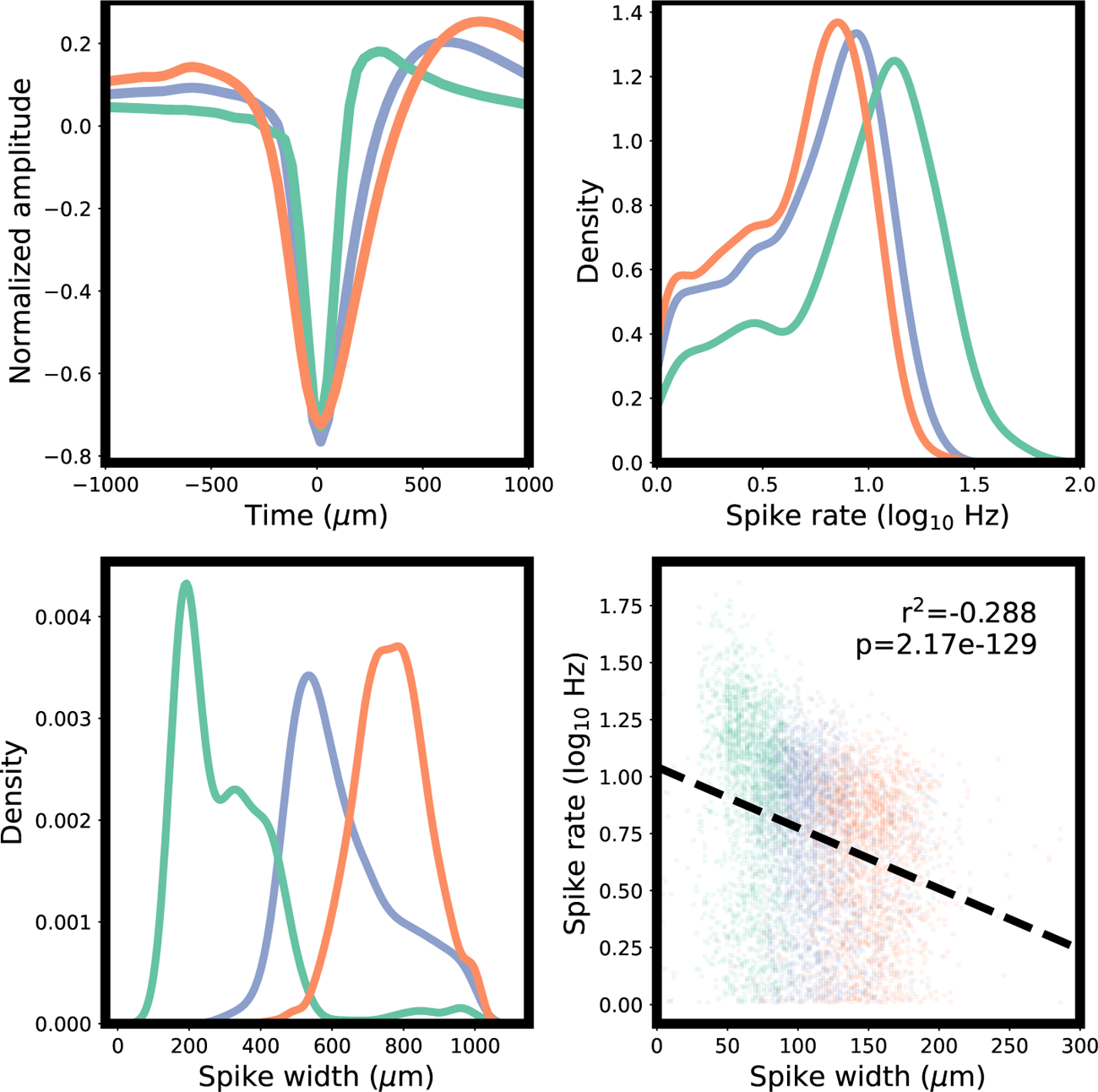
Spike widths and rates for each unit type. Spike width is given as the time between trough and peak.

Although the observed spike shape clusters were observed across all nuclei (Fig 11), spike shapes were distributed unequally (*χ*^2^(6,N=6754)=968.95, p*<*1e-5; Fig 11C-E).

**Figure 11:**
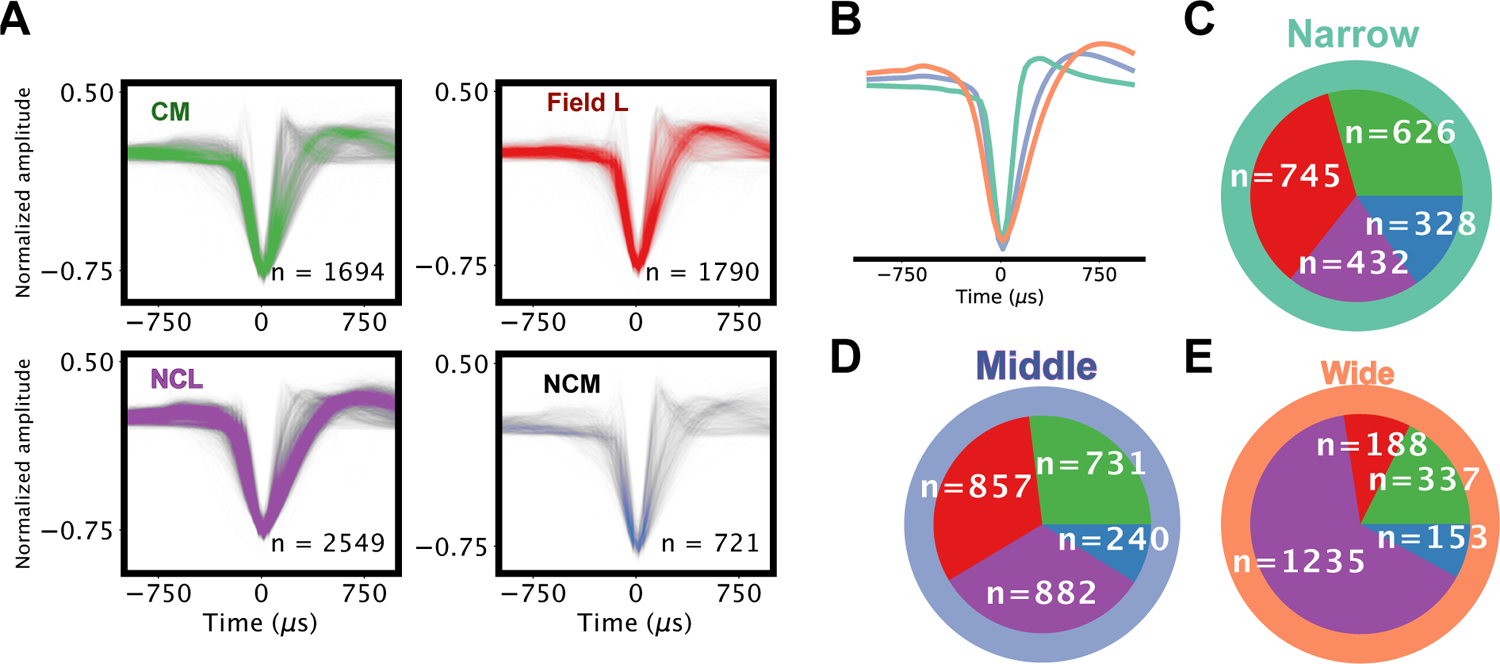
Spike shapes by unit locations. (A) Amplitude-normalized voltage traces of peak channel activity for all categorical units (see Methods 4.24) used in the analysis for each brain region. (B) Average trace of each unit-type cluster (C-E) Brain regions for each unit type.

### 4.21 Neural feature representation and response similarity

We represented spike trains as vectors using the methods outlined in Fig 3A-F). In particular, a PSTH of spike trains was computed with 10ms time-bins, which was then smoothed with a Gaussian kernel with a *σ* of 25ms. Morphs were sampled at a resolution of 128 points. For physiological analyses, we reduced the sampling resolution, binning the 128 interpolation points into 16 points along the morph, thus the neural response vectors and similarity matrices are 100 time-bins by 16 interpolation bins, and 16 interpolation bins by 16 interpolation bins, respectively.

We computed neural response similarity as the cosine similarity of the Gaussian convolved spike vectors, which has been effectively used to find similarity in spike trains in the past (*38*). A number of different similarity metrics could have been used in its place, for example, correlation coefficients (*39, 40*) and Euclidean distance between Gaussian convolved spike trains. We compared the cosine similarity to several other similarity metrics used in neural analyses including the correlation coefficient, Euclidean distance, and Manhattan distance, and found broadly similar results (Fig 12).

**Figure 12:**
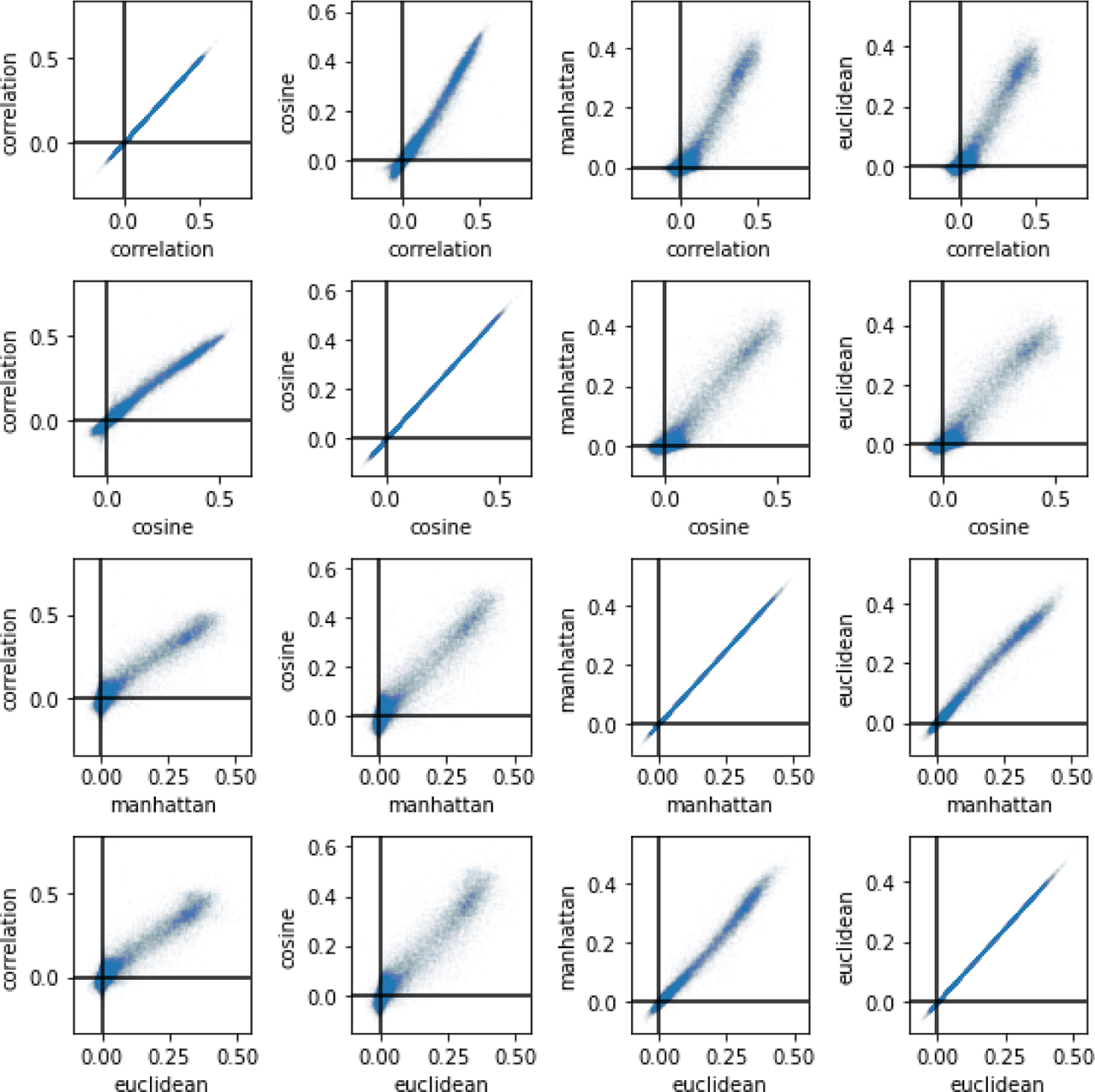
Comparison of unit categoricality using cosine similarity, correlation coefficient, Euclidean distance (1*/*(1 + *D*)) and Manhattan distance (1*/*(1 + *D*)).

### 4.22 Estimating a neurometric function from the similarity matrix

The neurometric function is computed on the basis of the similarity matrix and is detailed in Fig 13. For each interpolation point, we took the average of the within and between-category similarity (*S_C_*_1_ and *S_C_*_2_) took the ratio 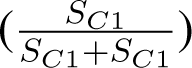 as the categorical similarity ratio. We then fit the same four-parameter logistic function as used in the psychometric function to the categorical similarity ratio as a function of interpolation point.

**Figure 13:**
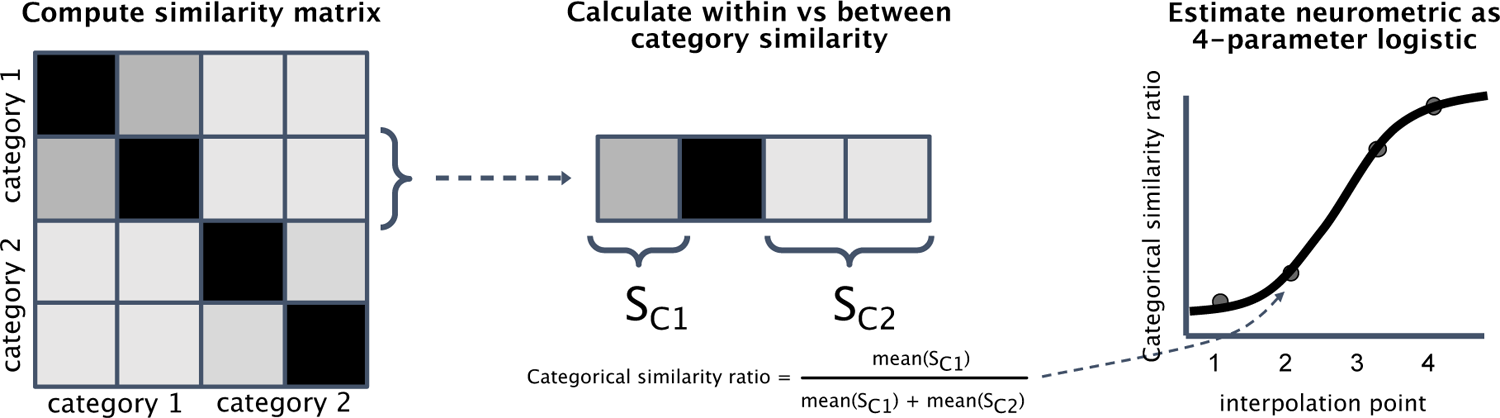
Method for computing a neurometric function from a similarity matrix.

### 4.23 Categoricality metric

Unit categoricality was computed using the similarity matrix (as seen in Fig 3). The similarity matrix used to compute a unit’s categoricality was the mean cosine similarity matrix across interpolation responses, where the cosine similarity matrix was computed over average response vectors for each interpolation point.

Similarity matrices were divided into four quadrants, corresponding to the within-category similarities for each category, and the between-category similarities. Categoricality was computed as the mean similarity in the within-category quadrants of the similarity matrix (i.e. the top left and bottom right), minus the between category similarities.

### 4.24 Subsetting categorical units

We operationalized behaviorally relevant, categorical, units on the basis of their response characteristics to the morph stimuli. Categorical units were determined by a threshold set in the categoricality metric. This threshold was set at a categoricality metric value above 0.1. These thresholds were set based upon visual assessment of unit responses (Fig 14) and similarity matrices.

**Figure 14:**
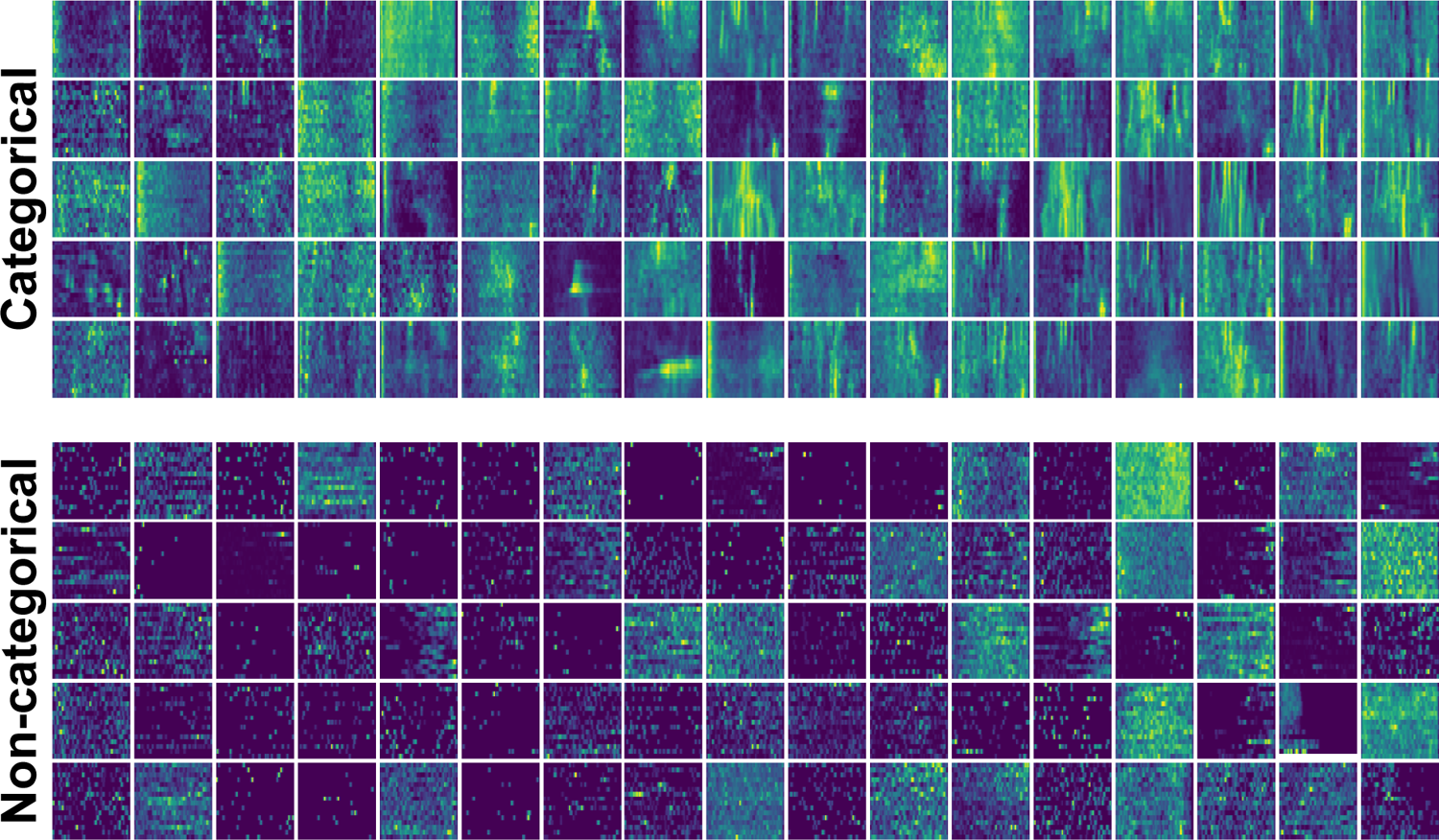
Categorical and non-categorical units, sorted by categoricality (right is greater).

### 4.25 Comparing spike rate across units, cues, and morphs

The analyses performed in section 2.2 were performed over unit spike rates in response to the morph stimuli, where spike rate was z-scored over the unit’s spike rates across all stimuli. A figure visualizing the main effect of cue and interactions between cue probability and stimulus class is shown in Figure 15. In addition, we shuffled the cue labels to ensure that our results were not due to inherent sampling biases present in the data (e.g. a left cue is more predictive of a left morph point, thus more cue left to left morph point samples exist in the dataset).

**Figure 15:**
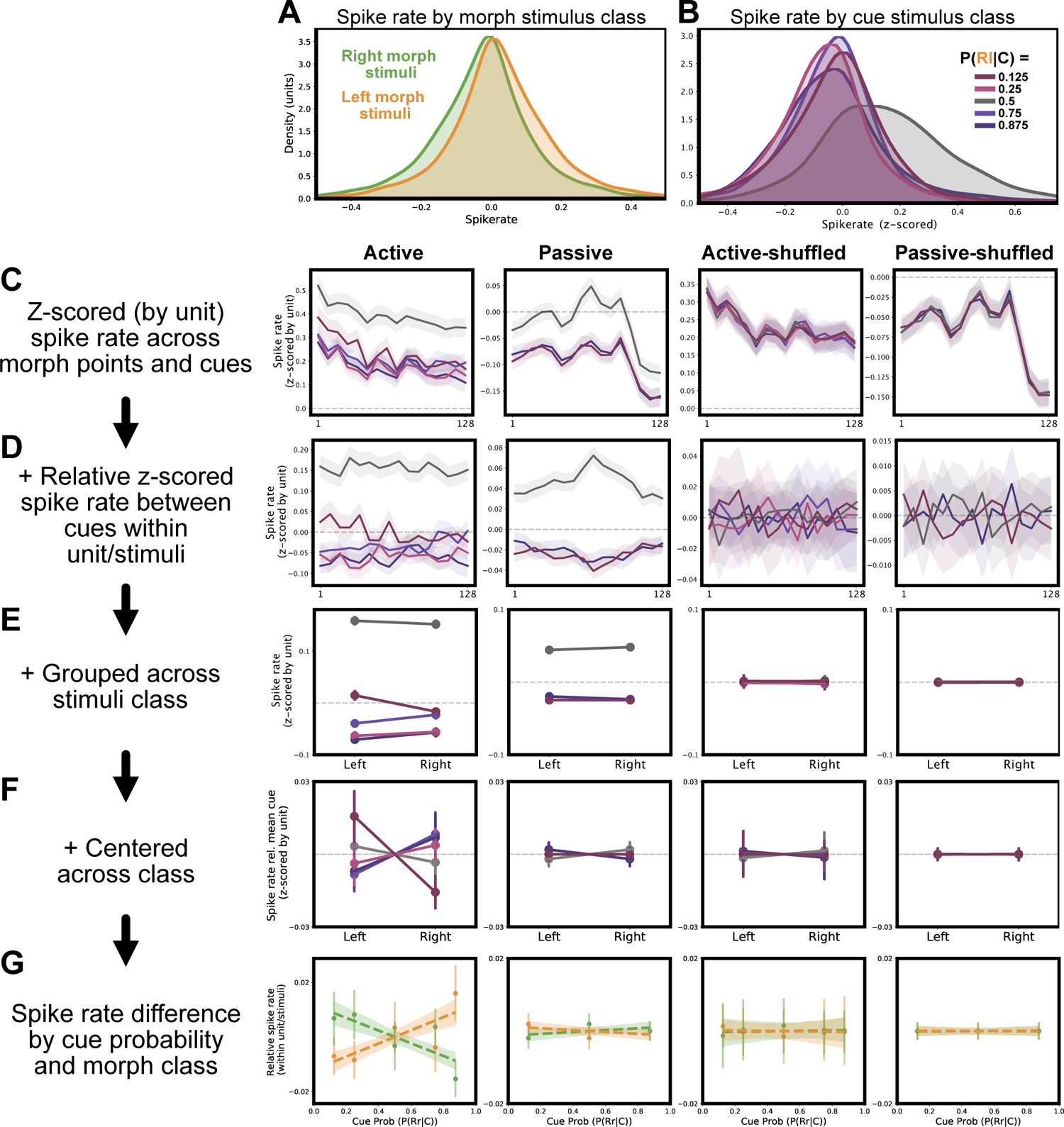
Spike rate modulation by cue. (A) Spike rate differs between stimuli. On average, across morphs, left stimuli elicit a greater spike rate than right stimuli. (B) Within stimuli, cues elicit differing spike rates for the same morph stimuli. Uncued trials yield the highest spike rate. (C) Across morph stimuli, average spike rates are shown for each cue condition in passive, and active conditions. Additionally, we perform the same analyses over data where cue labels are shuffled in the active and passive conditions (thus, no difference between spike rates across cues are observed). (D) The same data as the four panels from (C), subtracting the spike rate for each stimulus averaged across cue conditions. (E) The same data as in the four panels from (D), shown across morph sides (left and right) rather than morph interpolation points. (F) The same data as in (E) centered for each morph at zero,^5^to^2^ show interactions between cue conditions and morph stimuli classes. (G) The same data as in (G) plotted for each morph stimuli class as a function of cue probability, exhibiting the relationship between cue probability and spike rate for each cue class.

### 4.26 Morph class and cue interactions by subject, brain region, unit type, and morph

In Figure 16 we plot the interactions between cue probability and morph stimulus class, controlling for the overall unit’s spike rates to stimuli and a main effect of cue responses.

**Figure 16:**
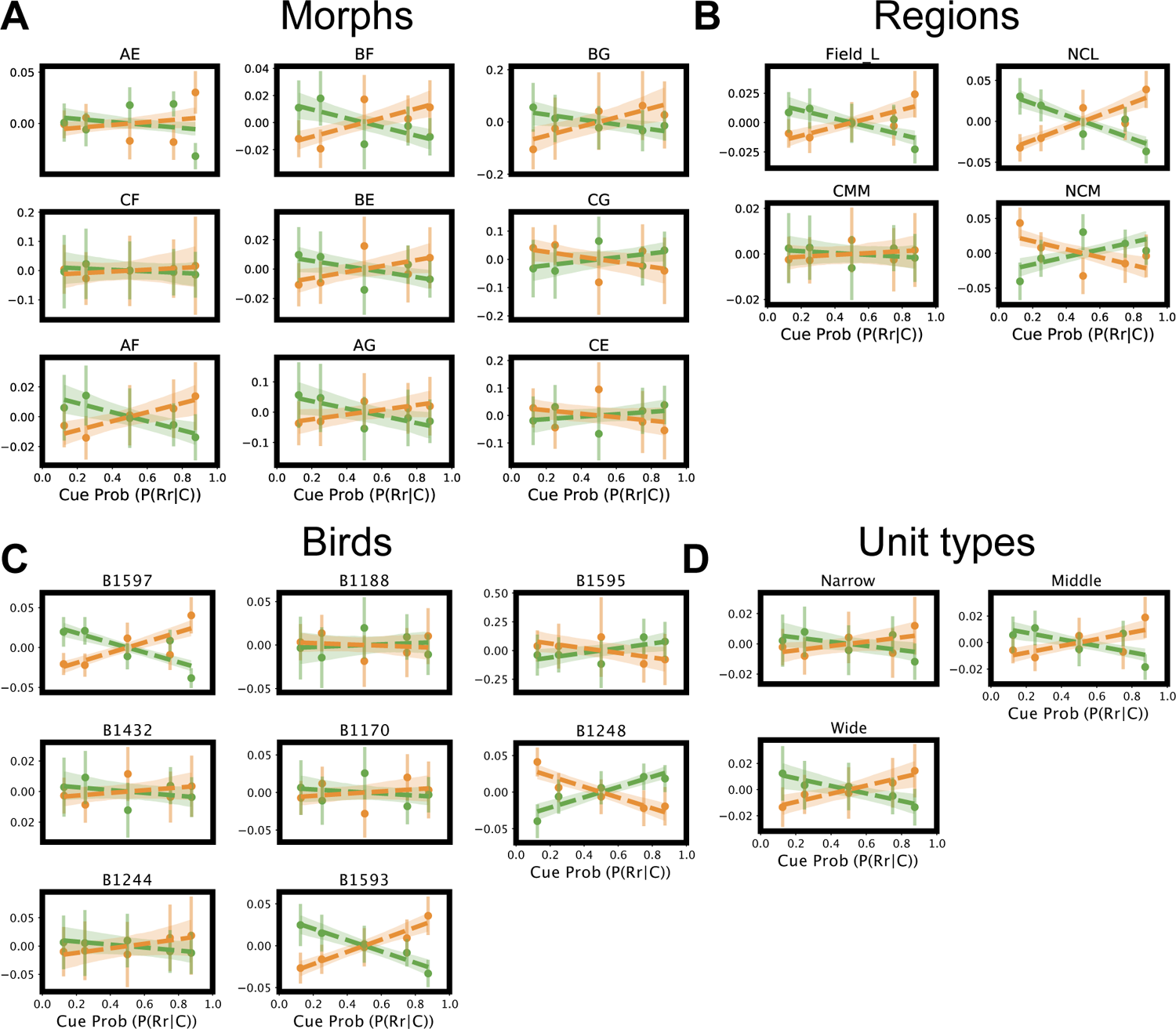
Interaction between cue probability and morph stimulus class on spike rate. Interactions are shown across four categories: Morph, brain region, subject, and unit class.

### 4.27 Differences in spike rate as a function of time

In section 2.2 we compared differences in spike rate as a function of their cue (i.e. within versus between cue).

To ensure that the effects of spike rate modulation occur between cue conditions, and not only between cue conditions and the uncued condition (where the main effect of cue on spike rate is greatest) we did not include the uncued condition in the spike rate differences between cue conditions in Fig 4B. We only included units and stimuli where we had active and passive behavioral trials (n units = 4722). We then, for each unit and stimulus, took the average absolute difference in response vectors between trials for trials with the same cue, and trials with different cues. The difference between the average absolute difference between cues, minus within cues, will equal zero when there is no difference between cue conditions. To ensure that no factors exogenous to between-cue differences are causing this effect, in Figure 17 we show the same analysis where cue labels have been shuffled within stimuli.

**Figure 17:**
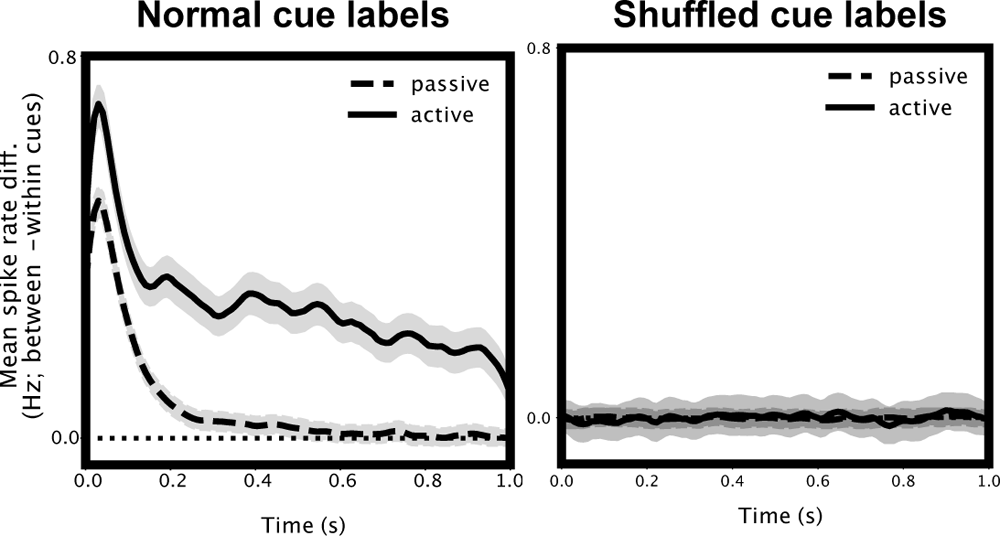
Spike rate differences between minus within cue categories over time. The right panel shows the same analysis with shuffled cue labels.

### 4.28 MNE receptive fields

We used the Maximum Noise Entropy (MNE) model to calculate receptive fields for all categorical units (*22, 41*) (4). MNE models were computed separately for each unit and predictive cue condition, resulting in four sets of receptive fields for each unit. During each model fitting, we used a jackknife procedure, averaging estimates from four subsets of the training data to yield the final parameters. For each unit on a given recording day, trials for each cue in which the expected target syllable category was presented (indicating a cue valid trial) were used to train the model. A number of valid trials equal to the number of invalid trials collected on the same recording day were held out from receptive field computation. These held-out trials were used to predict the receptive field model’s performance on both valid and invalid trials. Predicted spike trains were correlated with actual spike trains on held-out trials (4E). The correlation values were then averaged across trials for a given predictive cue, cue validity, and unit identity on an individual recording day to obtain a set of values for each unit corresponding to how well the receptive field model predicted valid and invalid trials in each cue condition (4F,G). This procedure was repeated four times with different random samples of held-out valid trials to ensure consistency of our results during fitting and analysis. The final results for each unit were averaged across these four samples. A linear mixed-effects model with a fixed effect for cue validity and random effects for each unit’s identity and recording day was used to predict trial correlation values. Paired t-tests were used for post-hoc testing of differences in cue valid versus cue invalid trials for each predictive cue.

### 4.29 Within cue response similarity

For each unit and cue, we computed the cosine similarity matrix across each morph. Cosine similarity matrices were computed by taking the average cosine similarity across trials for each interpolation point (16) in the morph. Analyses were only performed over active behavioral trials, where the subject provided a response. We then contrasted the cosine similarity matrices across different cue conditions. Figure 20 (top left) shows the average cosine similarity across left cues subtracted by the average cosine similarity across right cues. Blue in the top left of the plot (the orange bounding box) depicts less similarity in the predicted left class in left-cued trials. The reverse is true for the red in the bottom right. We measured this relationship showing that predicted morph classes are less similar within-class in Figure 20 (top right). Each point and confidence interval consists of the within-class similarity relative to the same unit’s response to uncued stimuli across trials. The negative relationship confirms that higher-probability cued trials exhibit less similar responses. In a similar manner as in Figures 15 and 17, we repeated this analysis over the same data in which cue labels had been shuffled within unit/interpolation. In the shuffled condition, we observe that the effect is removed. Finally, we broke out the analyses from Figure 20 in Figure 18 and 19, which are discussed in the main text.

**Figure 18:**
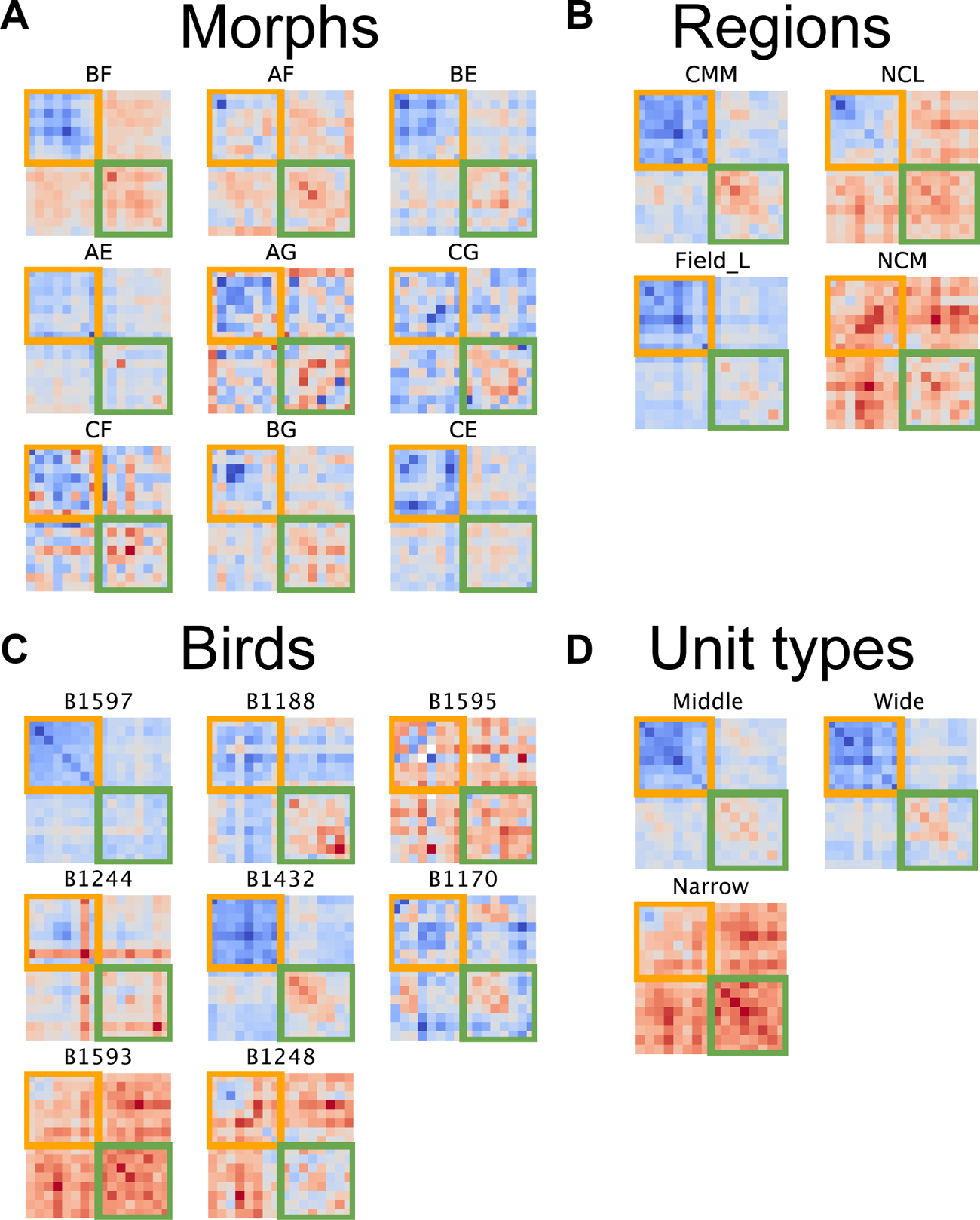
The similarity from Fig 5I broken out into individual morphs, brain regions, subjects, and unit types. The similarity matrix depicts the shift in spike train vector cosine similarity for left-cued minus right-cued trials. The shift is depicted here averaged is averaged across units.

**Figure 19:**
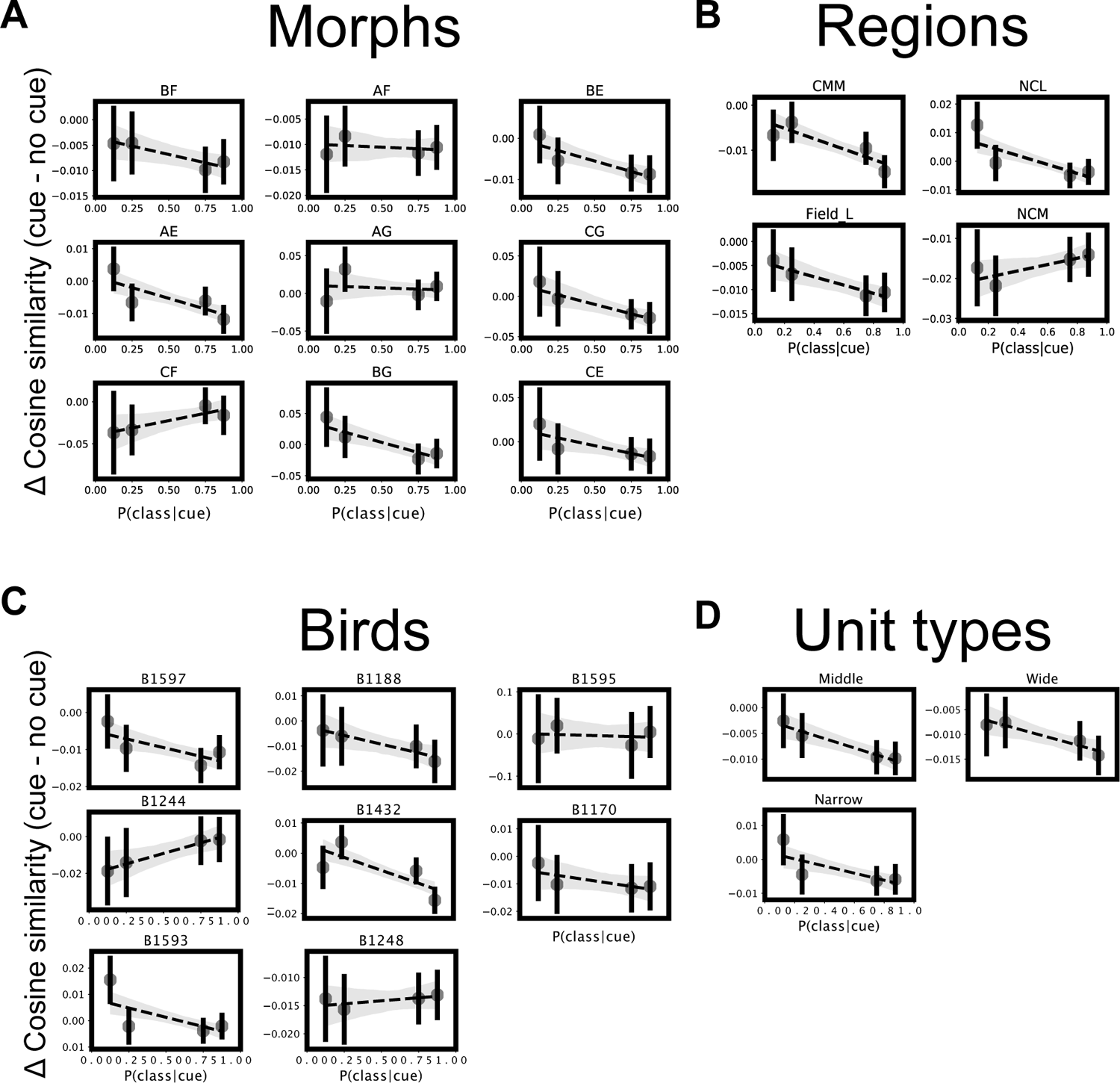
The relationship between the probability of the stimulus class and the shift in similarity from baseline (the uncued condition) shown in Fig 5J, broken out into morphs, brain regions, subjects, and unit types.

**Figure 20:**
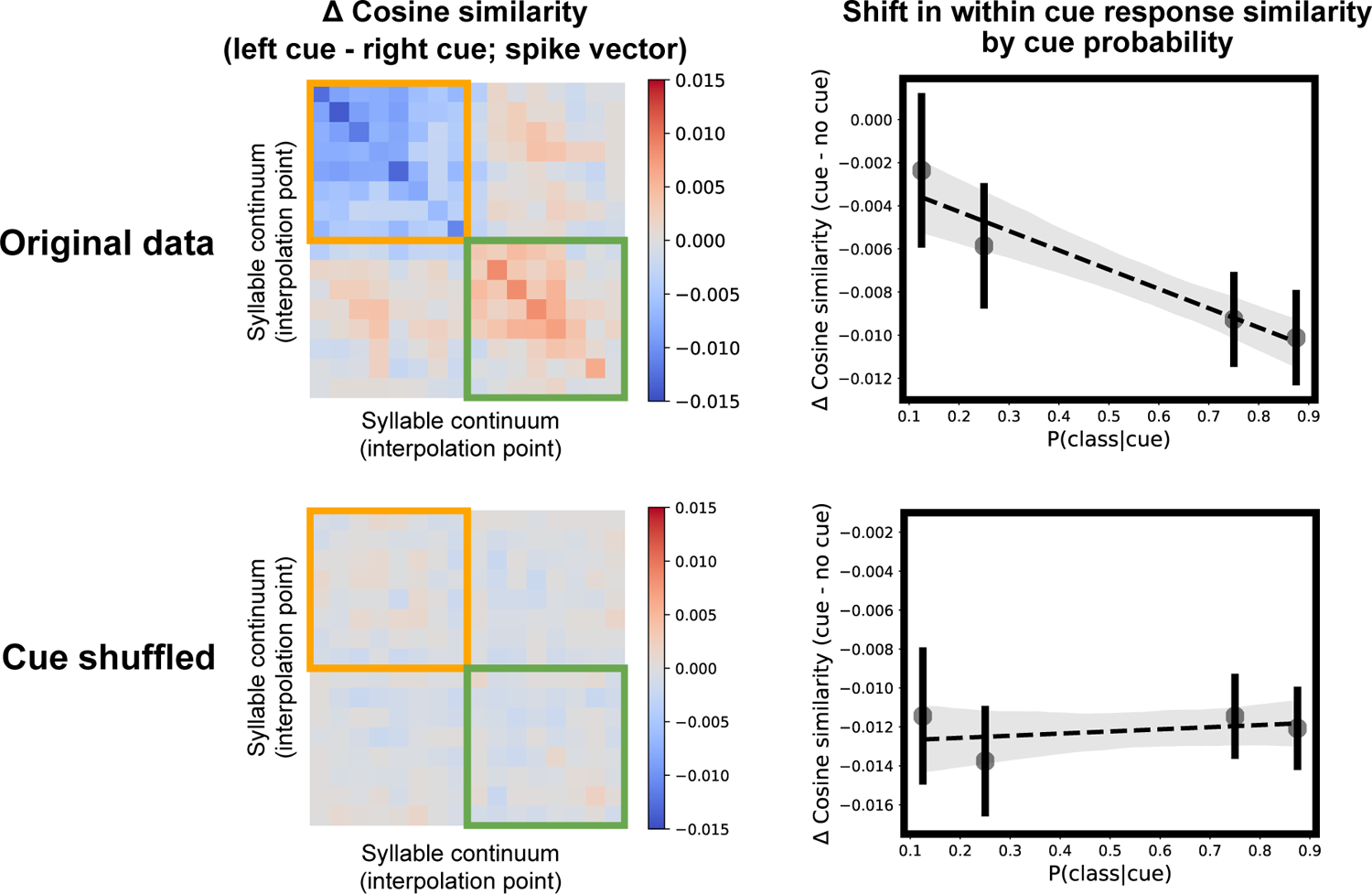
Spike vector cosine similarity and shift in similarity as a function of class probability as seen in Fig 5 I and J, compared with the same analysis performed over the same dataset where cue labels are shuffled.

